# Differential effects of the D1/S264V mutation in Photosystem II with either PsbA1 or PsbA3 on Q_B_, non-heme Iron, and the associated hydrogen-bond network

**DOI:** 10.1101/2025.05.14.654172

**Authors:** Kosuke Tada, Kaho Yamagata, Kazumi Koyama, Julien Sellés, Alain Boussac, Miwa Sugiura

## Abstract

The role of the D1/S264 residue and the role of its environment in the proton-coupled electron transfer reaction on the acceptor side of Photosystem II were investigated. To this end, D1/S264V mutants were constructed in the thermophilic cyanobacterium *Thermosynechococcus elongatus*, with D1 being either PsbA1 or PsbA3. The PSII mutants were investigated using EPR spectroscopy, thermoluminescence, (time-resolved) absorption changes measurements, and oximetry. While the mutation had minor effects in PsbA1-PSII, the S264V mutation in PsbA3-PSII had significant consequences: *i*) thermoluminescence data show inefficient electron transfer from QA^-^ to QB; *ii*) re-oxidation of QA^-^ was slowed, by at least a factor of 10; *iii*) the herbicides inhibit weakly O2 evolution; *iv*) no Fe^2+^QB^-^ EPR signal was detected in dark-adapted PSII; instead, *v*) a large Fe^3+^ signal was present with *vi*) modified EPR properties; *vii*) no QA^-^Fe^2+^QB^-^ biradical signal was observed after illumination at 198 K following a flash illumination, confirming the inefficient formation of QB^-^; *viii*) either no proton uptake coupled to non-heme iron reduction occurred or with a very slow rate compared to PsbA3-PSII; *ix*) changes were noted in the electrochromic response associated with QA^-^ formation; and *x*) increased production of singlet oxygen, both with and without herbicides. The S264V mutation in PsbA3-PSII leads to a significant decrease in the energy gap between the QA^-^QB and QAQB^-^ states. The effects listed above are discussed regarding the differences between PsbA1-PSII and PsbA3-PSII as those related to the sulfoquinovosyldiacylglycerol, the water molecules and the H-bond network.

## Introduction

Oxygenic photosynthesis in cyanobacteria, algae, and higher plants transforms solar energy into the chemical bonds of sugars and dioxygen [1]. Photosystem II (PSII) initiates this process by splitting water molecules to extract electrons, producing reduced quinones, generating a proton gradient, and releasing O2. The mature PSII complex contains 35 chlorophyll *a* molecules (Chl-*a*), 2 pheophytins (Phe-*a*), 1 membrane-bound b-type cytochrome (Cyt*b*559), 1 extrinsic c-type cytochrome in cyanobacteria and red algae (Cyt*c*550) 1 non-heme iron, 2 plastoquinones 9 (PQ9), QA and QB, the Mn4CaO5 cluster, 2 Cl⁻ ions, 12 carotenoids, and 25 lipids [2,3].

Among the 35 Chl-*a* molecules, 31 function as antenna chlorophylls. When one of these is excited, the excitation energy is transferred between chlorophylls until it reaches the key pigments in the photochemical reaction center constituted of 4 Chl-*a* molecules (PD1, PD2, ChlD1, and ChlD2) and 2 Phe-*a* molecules (PheD1 and PheD2), see Fig. 1A. A few picoseconds after the formation of excited *ChlD1, charge separation occurs, ultimately leading to the formation of the ChlD1^+^PheD1^-^ and then PD1^+^PheD1^-^ radical pair states [4,5]. The formation of the ChlD1^+^PheD1^-^ radical pair has been proposed as part of a “fast pathway” (short-range charge separation), in contrast to a “slow pathway” in which PD1PD2 acts as the initial donor (long-range charge separation), directly forming the PD1^+^PheD1^-^ radical pair [6].

**Figure 1:**
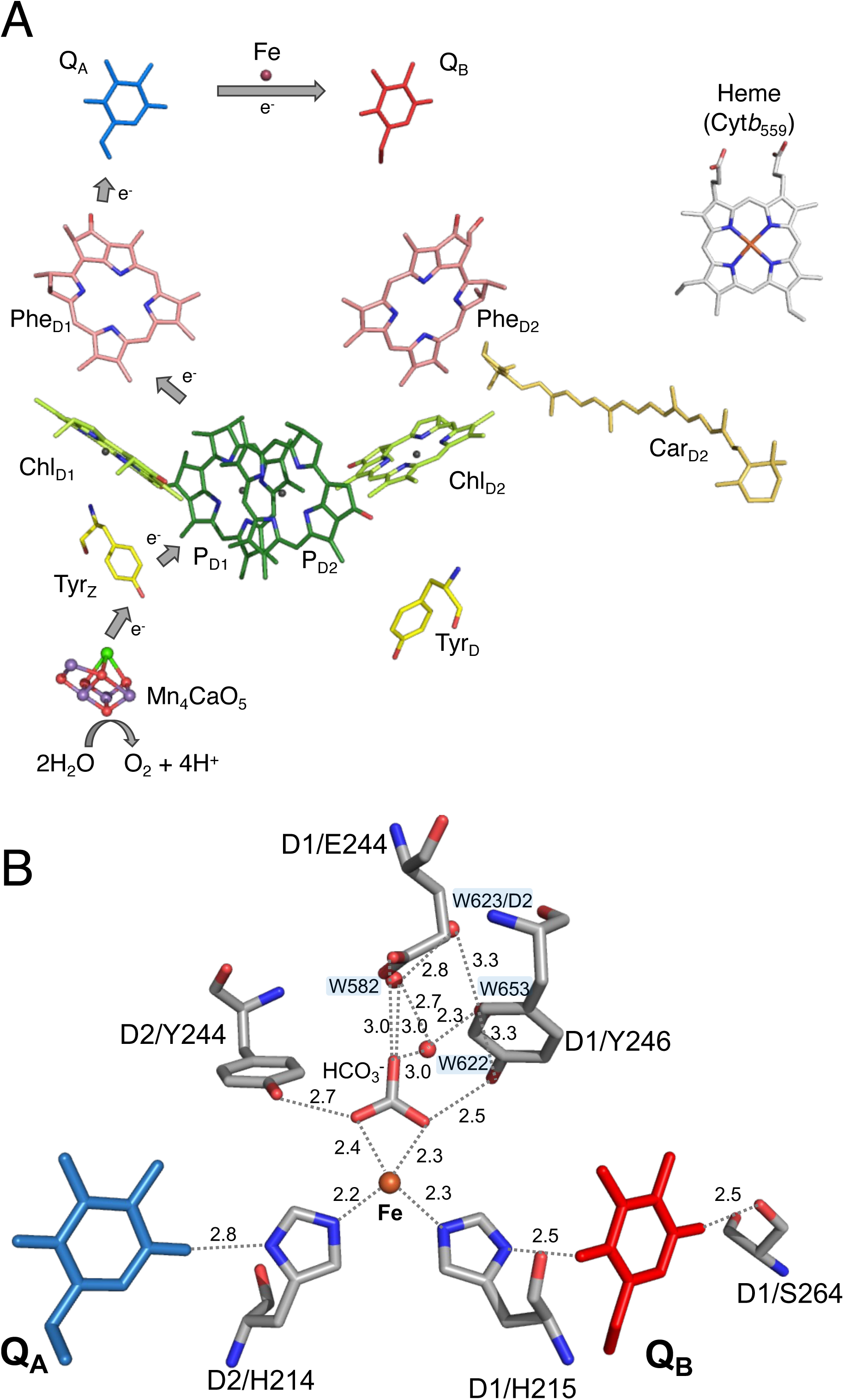
Arrangement of cofactors for electron transfer in PSII reaction center (A), structure around plastoquinone-binding sites including non-heme iron, bicarbonate and some water molecules (B), and structure of network between QA to QB (C). The figures were drawn with MacPyMOL with the A monomer of PsbA1-PSII in PDB 4UB6.

Following charge separation, PD1^+^ oxidizes TyrZ (Y161 of the D1 polypeptide), and then the oxidized TyrZ oxidizes the Mn4CaO5 cluster. The electron on PheD1^-^ is transferred to QA, the primary quinone electron acceptor, and then to QB, the secondary quinone electron acceptor (Fig. 1, Panel A). Under moderate light conditions, QA can only be singly reduced (QA^-^), whereas QB accepts two electrons and two protons. Then QBH2 leaves its binding site and is replaced by an oxidized QB molecule from the membrane plastoquinone (PQ) pool, reviewed in [7,8].

The Mn4CaO5 cluster, oxidized by the TyrZ^●^ radical generated after each charge separation, cycles through five redox states, denoted as Sn, where n represents the number of stored oxidizing equivalents. Upon formation of the S4-state, two water molecules bound to the cluster are oxidized, O2 is released, and the S0-state is reformed [9–13].

The electron transfer between QA and QB, along with the proton-coupled electron transfer (PCET) reactions involved in QB reduction [2,3,14], Panels A and B in Fig. 1, remain the focus of numerous experimental and computational studies. This process is a key aspect of the energetics of PSII, *e.g.*, [15–22].

Mutations of the D1/S264 amino acid have been shown to affect the resistance to herbicides which bind into, or very close to, the QB site and therefore block the electron transfer from QA^-^ to QB, *e.g.*, [23–29]. This resistance is consistent with structural changes in the QB binding pocket, where the distal oxygen of QB interacts with the side chain of D1/S264, along with D1/H252, potentially forming part of a proton transfer pathway to the reduced QB [16,22], Fig. 1B. More precisely, in computational works it has been proposed that: *i*) D1/S264 acts as a relay between D1/H252 and QB^-^ [16] and *ii*) the second proton transferred to QBH^-^ originates from D1/His215. The pathway for the reprotonation of D1/His215 may involve the bicarbonate bound to the non-heme iron, D1/Y246, and water molecules within the QB site [16], Fig. 1B. Beyond its role in proton transfer, molecular dynamics simulations suggest that D1/S264 undergoes reorientation during the release of QBH2 [17] and thus is involved in the QBH2/QB exchange [17,30].

In the presence of the artificial electron acceptor PPBQ, the non-heme iron (Fig. 1) is oxidized according to the reaction: Fe^2+^(QA/QB)^-^ + PPBQ → Fe^2+^(QA/QB) + PPBQ^-^→ Fe^3+^(QA/QB) + PPBQH2 [31]. In the cyanobacterium *Thermosynechococcus elongatus* the non-heme iron is oxidized in a small proportion of dark-adapted PSII [32] while PPBQ has not been added. In the absence of PPBQ, the oxidation of the non-heme iron is likely attributable to the equilibrium Fe^2+^ + QB^-^ ⇌ Fe^3+^ + QB^2-^ in a situation where QB^-^ becomes strongly oxidizing. Coupled to the oxidation of the non-heme iron, the release of a proton into the bulk is observed [33,34]. It has been proposed that the protonated D1/H215 residue is responsible for this proton release [33,34].

Including protons in the Fe^2+^ + QB^-^ ⇌ Fe^3+^ + QB^2-^ equilibrium highlights that the process involves a remodeling of the hydrogen-bond network around the Fe/QB site as the equilibrium is established. Conversely, alterations in the hydrogen-bond network, such as those resulting from the PsbA1/PsbA3 exchange [35,36], are expected to impact this equilibrium. The amount of Fe^3+^ has indeed been observed with a larger proportion in PsbA3-PSII than in PsbA1-PSII [37].

The role of the D1/S264 residue and the role of its environment in the proton-coupled electron transfer (PCET) reaction on the acceptor side have been further investigated in the present work. To this end, D1/S264V mutants were constructed in the thermophilic cyanobacterium *Thermosynechococcus elongatus*, with D1 being either PsbA1 or PsbA3. Several reasons motivated the construction of this mutant in both PsbA1 and PsbA3. Among them, the 21 amino acid differences between PsbA1 and PsbA3 result in significantly higher O2 evolution activity in PsbA3-PSII compared to PsbA1-PSII. This difference arises primarily from changes on the acceptor side, where the limiting step involves the QBH2/QB exchange. Notably, the substitution of S270 in PsbA1 with A270 in PsbA3 has been suggested to affect the stabilization of sulfoquinovosyldiacylglycerol (SQDG), which is positioned between QB and the non-heme iron in PsbA1 and PsbA3 [2,35]. Furthermore, the *E*_m_(QA/QA⁻) is shifted upward by approximately +41 mV in PsbA3-PSII compared to PsbA1-PSII [38], and the *E*_m_(PheD1/PheD1^-^) is higher by approximately +17 mV in PsbA3-PSII compared to PsbA1-PSII, which results in a larger energy gap (by 41-17=24 mV) between Phe Q ^-^ and PheD1^-^QA in PsbA3-PSII [39] (see Fig. S10 for a simplified energetic scheme). The PSII mutants were investigated using EPR spectroscopy, thermoluminescence (TL), (time-resolved) absorption changes measurements, and oximetry.

Among the main findings, it was observed that the S264V mutation with D1 = PsbA1 had minimal impact on the properties of the PSII acceptor side. In contrast, the S264V mutation with D1 = PsbA3 induced significant changes, including; modification of the Fe^2+^ + QB^-^ ⇌ Fe^3+^ + QBH2 equilibrium, a modified (much slower or lacking) proton uptake associated with non-heme iron reduction, a large decrease in the energy gap between the Q ^-^ QB and QAQB^-^ states, a higher singlet oxygen production, and a modified QB/herbicide binding site. These effects are discussed regarding the differences between PsbA1-PSII and PsbA3-PSII including the sulfoquinovosyldiacylglycerol (SQDG), the water molecules and the H-bond network.

## Material and Methods

### Construction of the mutants and purification of the PSII

The PsbA1/S264V and PsbA3/S264V mutants were constructed using the protocol described in Fig. 2. Site-directed mutations were introduced in the *psbA1* and *psbA3* genes to replace D1/S264 with D1/V264. The modification was made around position +790 of *psbA1* and *psbA3* on the corresponding plasmids using the QuickChange Lightning Site-Directed Mutagenesis Kit (Agilent Technologies), as illustrated in Fig. 2A. The segregation of all *psbA3* or *psbA1* copies in the genome of the deletion mutant was confirmed by PCR amplification of the mutation region, followed by *Bts* I-v2 digestion of the PCR products. The primers used were:

- Forward primer: 5’–CTACAACGGTGGCCCCTACCAACTGATCAT–3’
- Reverse primer: 5’–AAGTCGAGGGGGAAGTTGTGAGCATTGCG–3’

**Figure 2:**
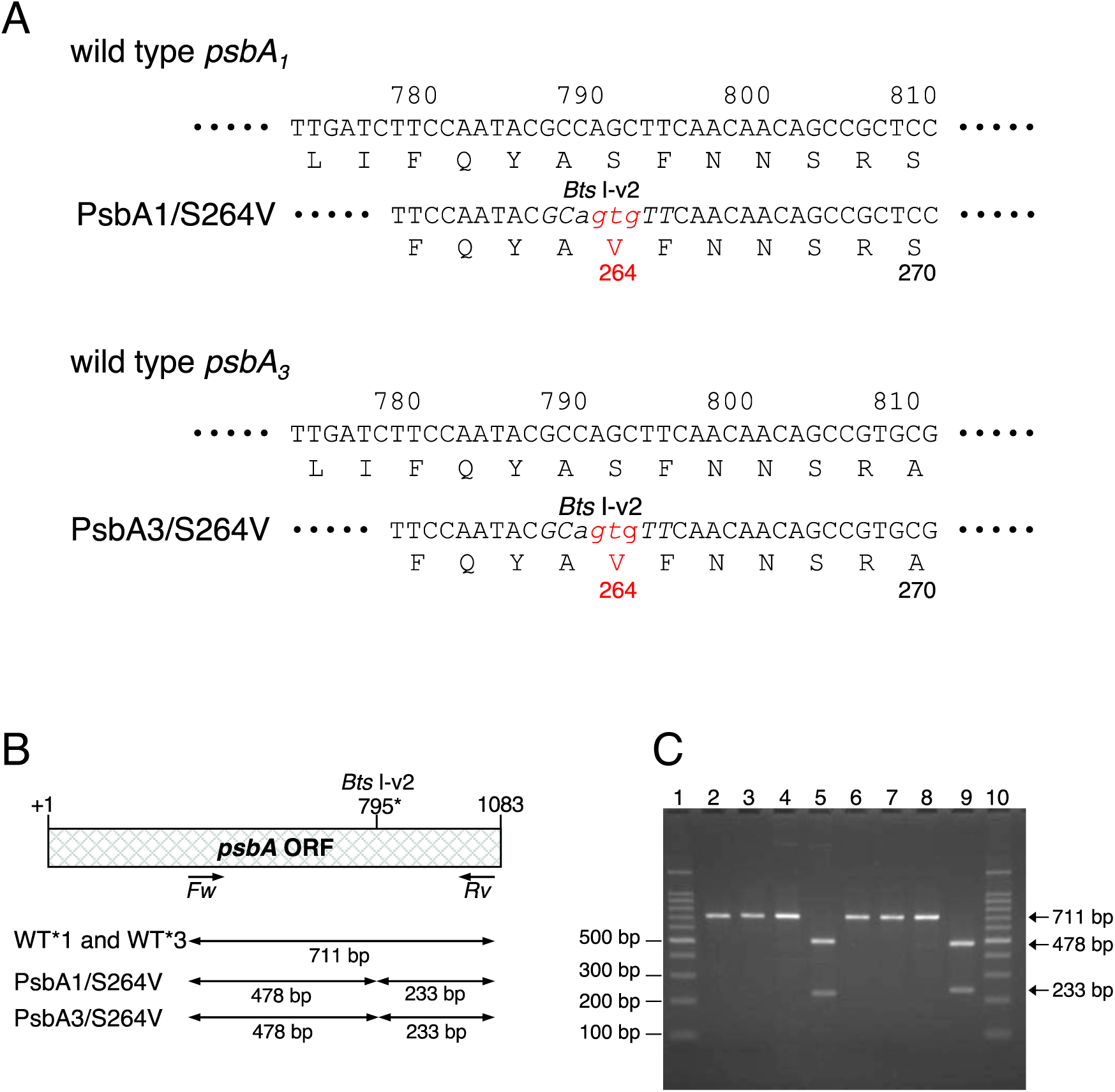
Nucleotide and amino acid sequences around D1/S264 to D1/V264 (A) and confirmation of the site-directed mutations of *T. elongatus* genome DNA (B, C). (A) Nucleotide sequence of *psbA1* and *psbA3* around PsbA1/S264 and PsbA3/S264. Positions of S264 are indicated with red letters. Numbers correspond to the position from the initial codon. Substituted nucleotides for S264V in both mutants are indicated in small letters. For making and selection of mutants, the *Bts* I-v2 restriction sites (letters in italic) was created in both *psbA1* and *psbA3* in the same time. (B) Physical map around *psbA*, and theoretical length of amplified DNA by PCR using *Fw* and *Rv* primers. The created *Bts* I-v2 site for the S264V mutants, are at position +795. The DNA fragments of 711 bp was digested with *Bts* I-v2 into 478 bp and 233 bp (C). Agarose gel (2%) electrophoresis of PCR products using the *Fw* and *Rv* primers (lanes 2 to 9) and the DNA fragments of the products after digestion with *Bts* I-v2 (lanes 3, 5, 7 and 9). Lanes 1 and 10, 100 bp DNA ladder markers (Nacalai, Japan); lanes 2 and 3, WT*1 strain [70]; lanes 4 and 5, PsbA1/S264V strain; lanes 6 and 7, WT*3 strain [41]; lanes 8 and 9, PsbA3/S264V strain.

The results of the digestion are shown in Figs. 1B and 1C.

As host cells, *T. elongatus* deletion mutants were used, in which: *i*) A His6-tag was introduced at the C-terminus of CP43 [40], and *ii*) *psbA1-psbA2* (for PsbA1/S264V) and *psbA2-psbA3* (for PsbA3/S264V) were deleted. Transformation of *T. elongatus* cells was performed by electroporation (BioRad Gene Pulser), and segregated cells were selected as described in [41]. Fig. 2 presents the results of the site-directed mutagenesis.

Cells of all strains were photo-autotrophically cultivated, and PSII were purified as previously described [39].

### SDS-polyacrylamide gel electrophoresis

For the SDS-polyacrylamide gel electrophoresis purified PSII’s (8 µg Chl) suspended in 40 mM MES/NaOH (pH 6.5), 10 mM NaCl, 10 mM CaCl2, 10 mM MgCl2, 0.03% *n*-dodecyl-β-D-maltoside were solubilized with 2% lithium laurylsulfate, and then analyzed by an SDS-polyacrylamide 16–22% gradient gel electrophoresis containing 7.5 M urea as previously described [42].

### Time-resolved absorption change measurements and EPR

Time-resolved absorption change measurements were conducted using a lab-built spectrophotometer [43], which was slightly modified as detailed in [44]. EPR spectroscopy was done as described previously [45].

### Oxymetry

Oxygen evolution of PSII complexes was measured at 25°C by polarography using a Clark-type oxygen electrode (Hansatech) with saturating white light through infrared and water filters at a Chl concentration of 2 µg of Chl mL^-1^ in 40 mM MES buffer (pH 6.5) containing 15 mM CaCl2, 15 mM MgCl2, and 10 mM NaCl. A total of 0.5 mM dichloro-*p-* benzoquinone (2,6-DCBQ, dissolved in dimethyl sulfoxide) was added as an electron acceptor. The PSII activity was measured immediately after the addition of 2,6-DCBQ. When used, the inhibitors 3-(3,4-dichlorophenyl)-1, 1-dimethylurea (DCMU), 3,5-dibromo-4-hydroxybenzonitrile (bromoxynil), 5-bromo-6-methyl-3-(1-methylpropyl)-2,4(1H,3H)-pyrimidinedione (bromacil) or 6-Chloro-*N*-ethyl-*N’*-(1-methylethyl)-1,3,5-triazine (atrazine), were added into the cuvette with PSII before light illumination.

Singlet oxygen production was detected by histidine mediated oxygen uptake measurements under high-light illumination conditions (20,000 µmol photons m^-1^ s^-1^) at 25°C, as previously described in [45–48]. The activity was measured in 40 mM MES buffer (pH 6.5) containing 15 mM CaCl2, 15 mM MgCl2, 10 mM NaCl in the presence of 200 mM L-histidine and 25 µM DCMU at 2 µg Chl mL^-1^ by using a Clark-type oxygen electrode (Hansatech).

### Thermoluminescence

TL glow curves were measured with TL500 (Photo System Instrument). Purified PSII were diluted to 100 µg Chl mL^−1^ in 40 mM MES buffer (pH 6.5) containing 1 M Betaine, 15 mM MgCl2, 15 mM CaCl2, 10 mM NaCl and then dark-adapted for 1 h at room temperature. Cells were diluted to 300 µg Chl mL^−1^ in the same buffer and then dark-adapted for 1 h after cw-light illumination at 30 mW cm^-2^ (MSG3-1100S-SD; Moritex) for 1 min. The samples were illuminated at 0°C by using one or two saturating xenon flashes with an interval of 200 ms. One second after the flash(s) illumination, the samples were heated at the constant heating rate 0.6°C s^−1^ and TL emission was detected. When 3-(3,4-dichlorophenyl)-1, 1-dimethylurea (DCMU) was added into the dark–adapted samples, TL emission was detected after one flash given at -15°C.

### Light-induced difference absorption spectra at 77 K

Light-induced difference absorption spectra at 77 K were measured basically as in our previous papers [45,46] using a spectrophotometer U-3900H (Hitachi High Tech. Co., Japan) equipped with a cryostat Optistat CF (Oxford Instruments) for low temperature measurements. The PSII were suspended in 20 mM MES buffer (pH 6.5) containing 15 mM CaCl2, 15 mM MgCl2, 10 mM NaCl and 65% glycerol at concentration of 0.2 mg Chl mL^-1^, and then dark adapted for 60 min on ice. After addition of 1 mM PPBQ, the PSII samples were illuminated for 30 s with a ∼ 30 mW cm^-2^ cw light source (MSG3-1100S-SD; Moritex Co., Japan) to induce the formation of QA^-^ (the oxidized species in this experiment being mainly the Cyt*b*559 as in [45,46]). The wavelength accuracy was ± 0.2 nm, the spectral band-pass was 2 nm and the scan speed was 120 nm min^-1^.

## Results

The two S264V mutant cells with either PsbA1 or PsbA3 were able to grow photo-autotrophically with growing rates similar to the WT*1 and WT*3 cells after the lag phase. The structural integrity and the activity of the purified PSII cores were monitored by: *i*) performing SDS-polyacrylamide gel electrophoresis, *ii*) measuring O2 evolution under continuous saturating light, and *iii*) recording the multiline EPR spectra arising from the Mn4CaO5 cluster in the S2 state induced by one-flash illumination at room temperature.

SDS-polyacrylamide gel electrophoresis (Fig. S1) revealed no loss of subunits in PSII from either mutant when compared to PsbA3-PSII.

Table 1 shows the oxygen evolution activity in the presence of 2,6-DCBQ, the added artificial electron acceptor. As reported previously [36], in the presence of 2,6-DCBQ, oxygen evolution in PsbA3-PSII , ∼ 5,000 µmol O2 (mg Chl)^-1^ h^-1^, was significantly higher than in PsbA1-PSII. However, while the oxygen evolution in PsbA1/S264V-PSII was only slightly reduced compared to PsbA1-PSII, the PsbA3/S264V mutant exhibited only 26% of the oxygen evolution activity observed in PsbA3-PSII. As demonstrated below by the similar amplitude of the S2 multiline signal recorded at 8.6 K in PsbA1/S264V-PSII and PsbA3/S264V-PSII (Figs. S3 to S5), the reduced O2 activity in PsbA3/S264V-PSII was not due to the loss of functional Mn4CaO5 but rather to the mutation’s impact on the acceptor side.

**Table 1:** Oxygen-evolving activity with the electron acceptor DCBQ and with and without herbicides

| Strain | no herbicides | +DCMU<sup>a)</sup> | +bromoxynil<sup>b)</sup> | +bromacil<sup>c)</sup> | +atrazine<sup>d)</sup> |
| --- | --- | --- | --- | --- | --- |
| WT*1 | 3,000 | 250 (92%) | 305 (88%) | 254 (92%) | 276 (91%) |
| PsbA1/S264V | 2,900 | 260 (91%) | 322 (89%) | 237 (92%) | 215 (93%) |
| WT*3 | 5,000 | 250 (95%) | 405 (92%) | 267 (95%) | 295 (94%) |
| PsbA3/S264V | 1,300 | 800 (38%) | 433 (67%) | 752 (42%) | 641 (51%) |

Unit of oxygen-evolving activity; $\mu\text{mol O}_2 (\text{mg Chl})^{-1} \text{h}^{-1}$ . The percentages are *versus* no herbicide. WT\*1 means PSII with wild-type PsbA1 and WT\*3 means PSII with wild-type PsbA3.
a) 25 $\mu\text{M}$ , b) 100 $\mu\text{M}$ , c) 100 $\mu\text{M}$ , d) 100 $\mu\text{M}$

The properties of QB, and more generally those of the acceptor side, in the PsbA1/S264V and PsbA3/S264V mutants were therefore further investigated using, firstly, EPR spectroscopy. Here, we summarize only the most significant results related to the acceptor side, specifically the spectra recorded at 4.2 K. All spectra recorded under other temperature conditions (8.6 K, and 15 K) are provided in the supplementary material (Figs. S3 to S6

In the first EPR experiment, no artificial electron acceptor was added. Panels A and B in Fig. 3 show the spectra recorded at 4.2 K for the PsbA1/S264V-PSII and PsbA3/S264V-PSII, respectively. Black spectra were recorded from PSII samples dark-adapted for 1 hour at room temperature. Red spectra were recorded after a single saturating laser flash at room temperature. Blue spectra were recorded after a further continuous illumination at 198 K following the flash illumination. In both samples, a bump between 0 and 500 gauss was observed, caused by contamination in the cryostat. However, this does not interfere with the results, as it completely disappears in the difference spectra, as shown in Panel C of Fig. 3.

**Figure 3:**
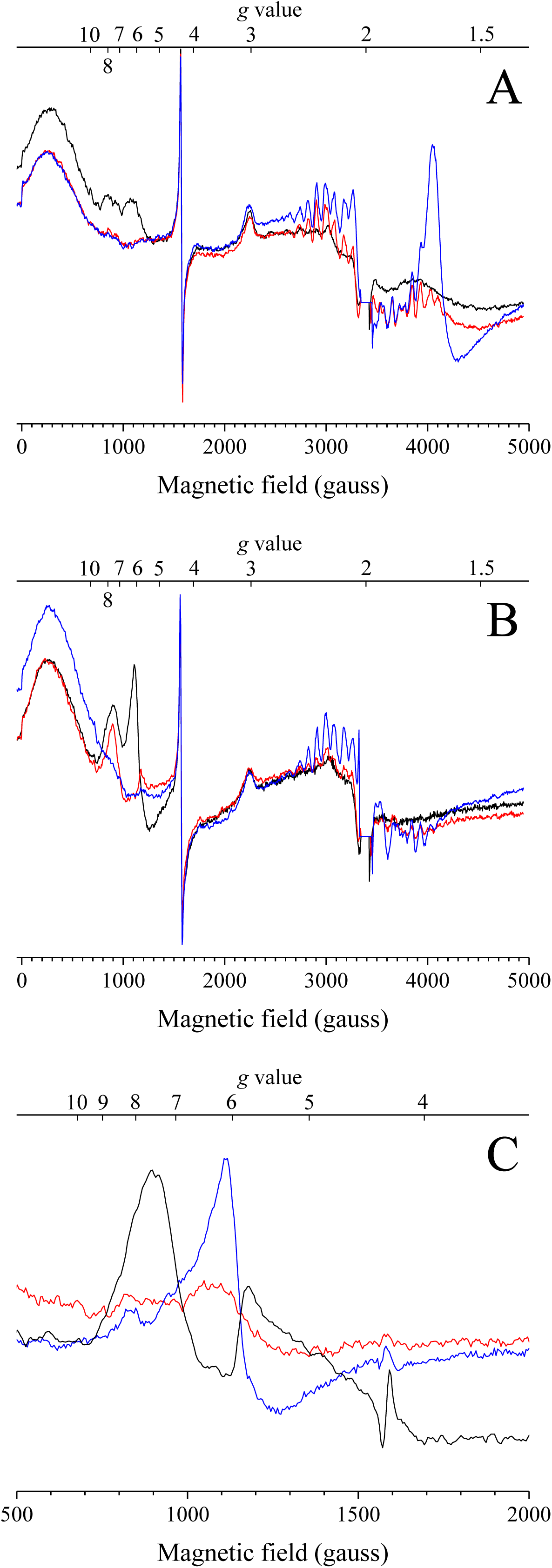
cw-EPR spectra recorded at 4.2 K in active PsbA1/S264V-PSII (Panel A) and active PsbA3/S264V-PSII (Panel B). Spectra in black were recorded in the dark-adapted samples. Spectra in red were recorded after laser flash illumination. The blue spectra were recorded after a further continuous white light illumination for 5-10 s at 198 K. Panel C shows difference spectra in the magnetic field region corresponding to the oxidized non-heme iron. The red spectrum corresponds to the “dark-*minus*-1 flash” difference in PsbA1/S264V-PSII. The blue spectrum corresponds to the “dark-*minus*-1 flash” difference in PsbA3/S264V-PSII. For comparison, the black spectrum corresponds to the “+PPBQ-*minus*-no add” difference in PsbA3-PSII. The microwave power was 20 mW. The modulation frequency was 100 kHz, the modulation amplitude was 25 gauss, and the microwave frequency was 9.49 GHz.

We will first describe the data for PsbA1/S264V-PSII (Panel A in Fig. 3), and two observations can be made. Firstly, the spectrum recorded at 4.2 K in dark-adapted PsbA1/S264V-PSII closely resembles that of PsbA1-PSII [49], with the exception that, by comparison with the other signals detected in the dark-adapted state, the amplitude of the Fe^2+^QB^-^ signal, seen at ∼ 4154 gauss and corresponding to *g* ∼ 1.63, appeared to be smaller than previously reported in the literature for the wild type PSII [37]. Secondly, the EPR signals from the oxidized non-heme iron signal have a slightly larger amplitude in the PsbA1/S264V-PSII than in PsbA1-PSII. The oxidized non-heme iron exhibits signals at ∼ 872 gauss (*g* ∼ 7.77) and ∼ 1121 gauss (*g* ∼ 6.05). These two observations suggest that the Fe^2+^QB^-^ ⇌^…^⇌ Fe^3+^QB equilibrium is slightly shifted to the right in PsbA1/S264V-PSII compared to PsbA1-PSII (see the Discussion section on this point).

After the single flash given at room temperature Q_A_^-^ is formed and the Fe^3+^ is then reduced by this Q_A_^-^ (red spectrum in Panel A of Fig. 3). This happens because the electron transfer from QA^-^ to Fe^3+^ is at least 10 times faster than the electron transfer to QB. As there is a small Fe^2+^QB^-^ signal in the dark, and therefore a large proportion of centers with oxidized QB, we could expect a large Fe^2+^QB^-^ signal after the flash illumination. This was however not observed because that was the Fe^3+^ present in the dark prior to the illumination that was reduced by the light-induced Q_A_^-^. We will come back on this point after analyzing the TL data.

After a further continuous illumination at 198 K (blue spectrum), the QA^-^Fe^2+^QB^-^ biradical signal seen at ∼ 4130 gauss which corresponds to *g* ∼ 1.64, see for example [20], was formed in centers in the QAFe^2+^QB^-^ state before the illumination at 198 K. The QA^-^Fe^2+^QB^-^ state is formed because the electron transfer from QA^-^ to QB^-^ is inhibited at 198 K [20].

In PsbA3/S264V-PSII (Panel B in Fig. 3), two main observations can be made from the spectrum recorded in the dark-adapted state (black spectrum): *i*) the absence of the Fe^2+^QB^-^ signal, and *ii*) a large non-heme iron signal with resonances at ∼ 917 gauss (*g* ∼ 7.39) and ∼ 1117 gauss (*g* ∼ 6.07). After a single flash at room temperature (red spectrum), the Fe^3+^ signal decreased significantly, leaving only the resonance at ∼ 917 gauss (*g* ∼ 7.39). The absence of a Fe^2+^QB^-^ signal after flash illumination suggests that either QB was missing, and/or the non-heme iron was oxidized in nearly all centers in the dark-adapted state since in this case, the electron on QA^-^ is expected to be transferred to the Fe^3+^ and not to QB. After the further illumination at 198 K (blue spectrum), a small QA^-^Fe^2+^ signal at ∼ 3570 gauss (*g* ∼ 1.9) was formed and the signal at ∼ 917 gauss (*g* ∼ 7.39) disappeared. The QA^-^Fe^2+^QB^-^ biradical signal at *g* ∼ 1.6 was also not induced by the further illumination at 198 K, thus supporting the conclusion that there was no Fe^2+^QB^-^ after the flash illumination.

Panel C in Fig. 3 shows the difference spectra in the magnetic field region corresponding to the oxidized non-heme iron. The red spectrum corresponds to the “dark-*minus*-1 flash” difference in PsbA1/S264V-PSII. The blue spectrum corresponds to the “dark-*minus*-1 flash” difference in PsbA3/S264V-PSII. For comparison, the black spectrum corresponds to the “+PPBQ-*minus*-no add” difference in PsbA3-PSII [31] with resonances detected at ∼ 922 gauss (*g* ∼ 7.5) and ∼ 1161 gauss (*g* ∼ 5.7). This non-heme iron signal in PsbA3-PSII in the presence of PPBQ was formed following the reactions Fe^2+^QB^-^ + PPBQ → Fe^2+^QB + PPBQ^-^ → Fe^3+^QB + PPBQH2 [31,50,51].

The shape of the three difference spectra in Panel C of Fig. 3 differ, indicating a change in the geometry of the Fe^3+^ orbitals among the three samples. The “dark-*minus*-1 flash” difference spectrum in PsbA1/S264V-PSII, while slightly different, closely resembles the non-heme iron signal in PsbA3-PSII (which is identical to that in PsbA1-PSII, not shown). In contrast, the “dark-*minus*-1 flash” difference spectrum in PsbA3/S264V-PSII is significantly different from that in PsbA1/S264V-PSII, displaying a much more axial signal. A second observation in PsbA3/S264V-PSII, evident in Panel B of Fig. 3, is that the signal lost upon the flash illumination differs from the signal lost upon subsequent continuous illumination at 198 K. This indicates structural heterogeneity at the Fe^3+^ site in PsbA3/S264V-PSII, with two Fe^3+^ that are reducible at different temperatures.

More explanations on the other signals seen in the spectra are given in the supplementary material. Briefly, in both PsbA1/S264V-PSII and PsbA3/S264V-PSII, the S2 multiline signal induced by a single saturating laser flash at room temperature, and recorded at 8.6 K, was very large, with an amplitude similar in the two mutants, while the further illumination at 198 K, expected to induce the S2 multiline signal in centers which remained in S1 after the flash illumination, induced a very small additional multiline signal. This observation supports the conclusion that these PSII samples are fully intact with a miss parameter similar to that generally observed in wild-type PSII from *T. elongatus*, ∼ 10%, *e.g.*, [52]. Additionally, in both PsbA1/S264V-PSII and PsbA3/S264V-PSII, the Cyt*c*550 signal, better seen at 15 K with a lower microwave power (5 mW), exhibited the same large amplitude, further confirming the intactness of the samples. It is worth remembering that at 4.2 K, a microwave power equal to 20 mW is strongly saturating rendering the amplitudes of the Cyt*c*550 signal at this temperature meaningless.

In the second EPR experiment, reported in Fig. 4, PPBQ was added to the samples after dark-adaptation for 1 hour at room temperature. Panels A and B show the spectra recorded at 4.2 K for the PsbA1/S264V-PSII and PsbA3/S264V-PSII mutants, respectively. The black spectra were recorded in dark-adapted PSII. The red spectra were recorded after a single saturating laser flash at room temperature. The blue spectra were recorded after an additional continuous illumination at 198 K following the flash illumination.

**Figure 4:**
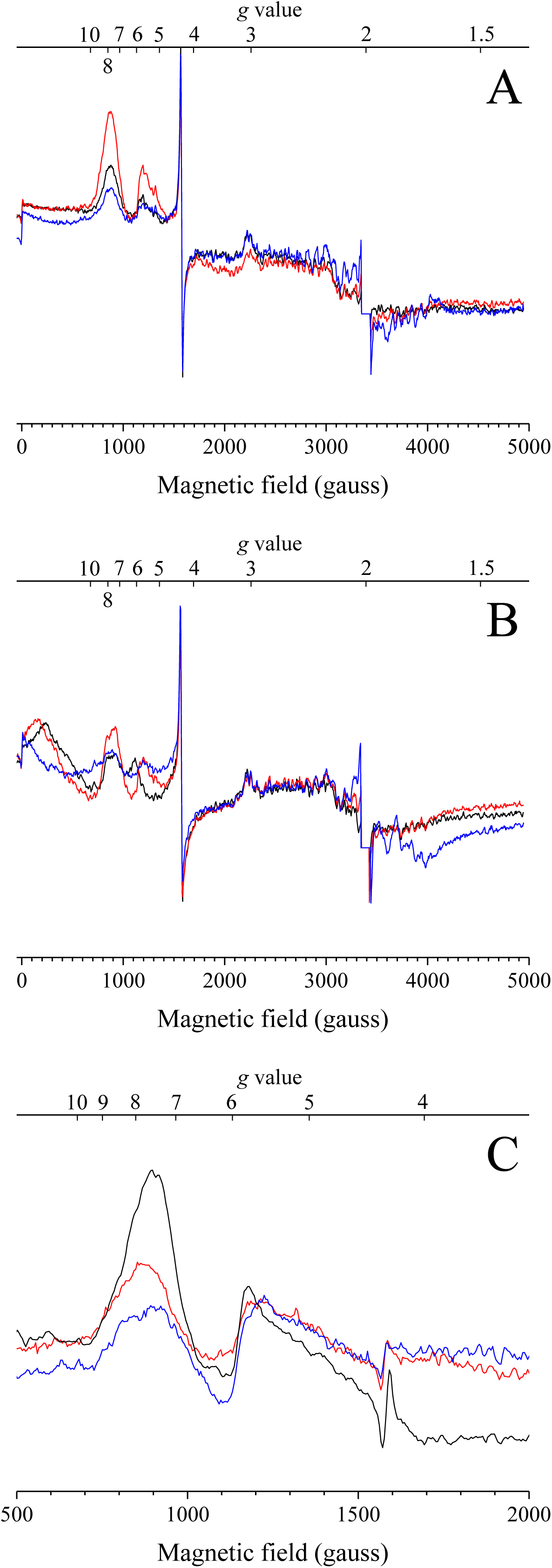
cw-EPR spectra recorded at 4.2 K, in active PsbA1/S264V-PSII (Panel A) and active PsbA3/S264V-PSII (Panel B). Spectra in black were recorded in the dark-adapted samples. Spectra in red were recorded after laser flash illumination. The blue spectra were recorded after a further continuous white light illumination for 5-10 s at 198 K. PPBQ dissolved in dimethyl sulfoxide was added to the dark-adapted samples prior to their freezing. Panel C shows difference spectra in the magnetic field region corresponding to the oxidized non-heme iron. The red spectrum corresponds to the “dark-*minus*-1 flash” difference in PsbA1/S264V-PSII. The blue spectrum corresponds to the “dark-*minus*-1 flash” difference in PsbA3/S264V-PSII. For comparison, the black spectrum corresponds to the “+PPBQ-*minus*-no add” difference in PsbA3-PSII. The microwave power was 20 mW. The modulation frequency was 100 kHz, the modulation amplitude was 25 gauss, and the microwave frequency was 9.49 GHz.

A comparison of these new spectra with those from the previous experiment conducted in the absence of PPBQ (Fig. 3) reveals differences in both the non-heme iron and quinone signals. In PsbA1/S264V-PSII, the non-heme iron signal is larger after a single flash due to its oxidation by the PPBQ^-^ that is formed in the PPBQ+(QA/QB)^-^ → PPBQ^-^+(QA/QB) reaction [31]. When compared to the experiment reported in Fig. 3, small differences in the non-heme iron signal may arise from the occupancy of the QB site by either QB or PPBQ [51]. After further continuous illumination at 198 K, two observations can be made: *i*) Fe^3+^ was largely reduced into Fe^2+^ (by the light-induced Q_A_^-^) in both samples, and *ii*) only a very small Q ^-^ Fe^2+^QB^-^ biradical and Q ^-^Fe^2+^ signals were detected in PsbA1/S264V-PSII. In PsbA3/S264V-PSII, the quinone signal induced by continuous illumination at 198 K was also very small, with no Q ^-^Fe^2+^QB^-^ biradical signal. Instead, a very small Q ^-^Fe^2+^ signal was observed, resembling the signal typically seen when no bicarbonate is bound to the non-heme iron, *e.g.*, [8].

Panel C in Fig. 4 shows difference spectra in the magnetic field region corresponding to the oxidized non-heme iron. The red spectrum corresponds to the “dark-*minus*-1 flash” difference in PsbA1/S264V-PSII. The blue spectrum corresponds to the “dark-*minus*-1 flash” difference in PsbA3/S264V-PSII. For comparison, the black spectrum corresponds to the “+PPBQ-*minus*-no add” difference in PsbA3-PSII. In the presence of PPBQ, the shape of the Fe^3+^ signals is similar in all three cases.

The consequences of the S264V mutation on the structural properties of the electron acceptors detected by EPR suggest that the energetics may also be affected. Such possible changes were investigated using TL. TL is a technic able to address the redox couples on both the donor side and the acceptor side. More precisely, concerning the acceptor side, TL addresses the energy gaps between PheD1^-^QA and PheD1QA^-^, and QA^-^QB and QAQB^-^ [53–55].

Fig. 5 shows the results of such a TL experiment. The heating rate was 0.6°C/s and the flash was given at -15°C and 0°C with and without DCMU, respectively.

**Figure 5:**
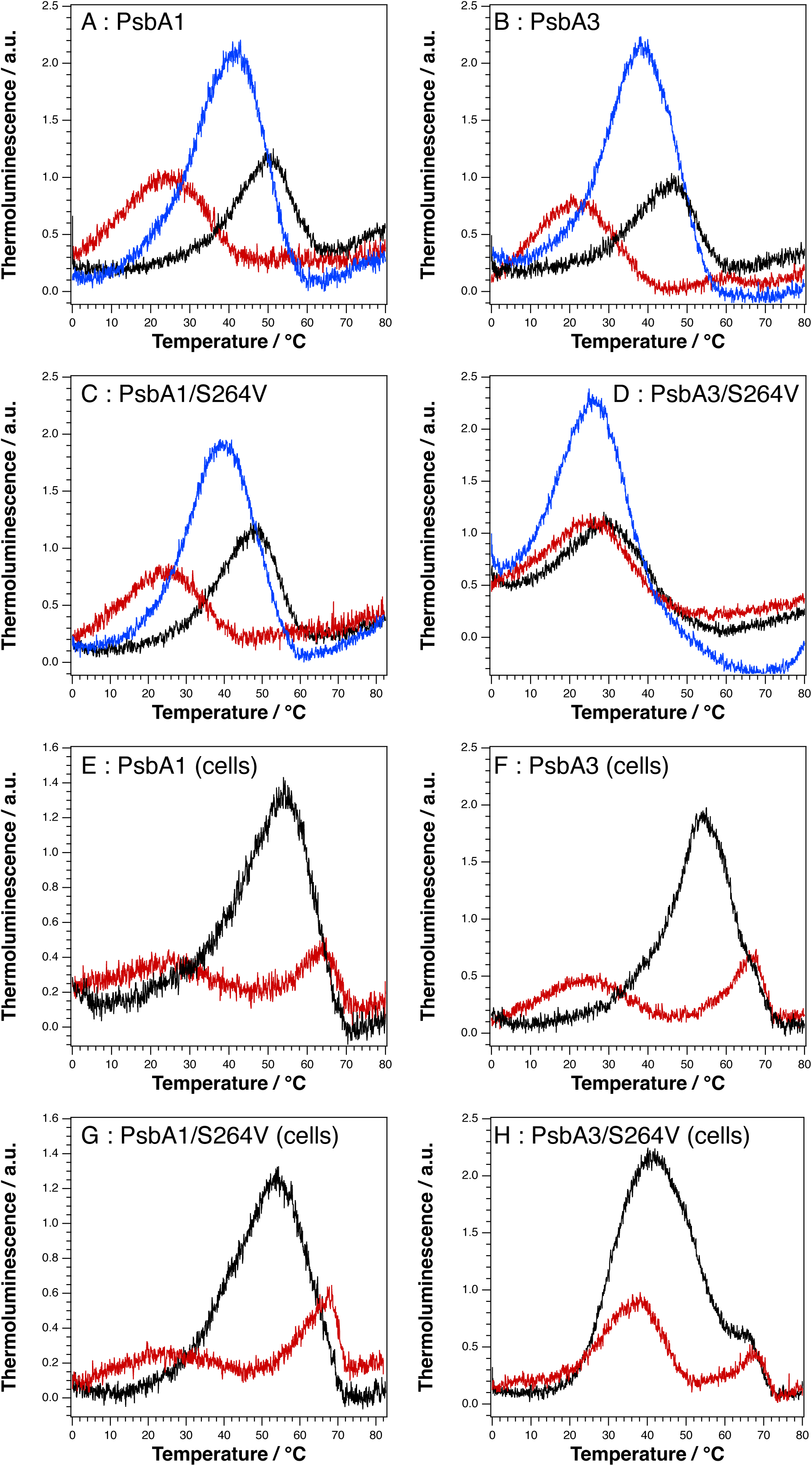
Thermoluminescence curves. Panel A was recorded in PsbA1-PSII, Panel B in PsbA3-PSII, Panel C in PsbA1/S264V-PSII, Panel D in PsbA3/S264V-PSII, Panel E in PsbA1-cells, Panel F in PsbA3-cells, Panel G in PsbA1/S264V-cells and Panel H in PsbA3/S264V-cells. The black curves correspond to TL emission recorded after 1 flash given in the absence of DCMU. The red curves correspond to a TL emission recorded after 1 flash given in the presence of DCMU. The blue curves were recorded after 2 flashes without DCMU. The samples were dark-adapted at room temperature for 1 h prior to the addition of DCMU. The heating rate was 0.6°C s^-1^.

Panel A was recorded in PsbA1-PSII, Panel B in PsbA3-PSII, Panel C in PsbA1/S264V-PSII, Panel D in PsbA3/S264V-PSII. Panels E and F were recorded in whole PsbA1-cells and PsbA3-cells, respectively. Panels G and H were recorded in whole PsbA1/S264V cells and PsbA3/S264V cells, respectively. The black curves correspond to the TL emission recorded after 1 flash given in the absence of DCMU. The red curves correspond to the TL emission recorded after 1 flash given in the presence of DCMU. The blue curves were recorded after 2 flashes in the absence of DCMU.

In the presence of DCMU (black curves), the TL arose from the S2QA^-^/DCMU charge recombination. In purified PSII, the amplitudes and peak temperatures at 24°C in PsbA1-PSII (Panel A), 22°C in PsbA3-PSII (Panel B), and 24°C in PsbA1/S264V-PSII (Panel C) do not greatly differ. At first glance, the difference between PsbA3-PSII and PsbA1-PSII appears smaller than expected. Nevertheless, this observation, discussed earlier in detail in [38,39], aligns with the experimental observations, the calculations and is consistent with the differences in ^1^O2 productions in PsbA1-PSII and PsbA3-PSII when all the recombination routes are considered that are the radiative routes (direct and indirect), the non-radiative route including the triplet route [39], and possible changes in the equilibria in the electron donor side [38].

In PsbA3/S264V-PSII (Panel D), the TL peak was very slightly shifted upward by ∼ 5°C and the amplitude was also slightly larger compared to PsbA3-PSII. A possible interpretation is a slightly larger energy gap between PheD1^-^QA/DCMU and PheD1QA^-^/DCMU in PsbA3/S264V-PSII. At this step there are two hypothesis including either a less reducing *E*_m_ for the QA/QA^-^ couple or a more negative *E*_m_ (PheD1/PheD1^-^) to account for a higher ^1^O2 production in this mutant (see Table 2). In whole cells, the peaks were at the same temperature, ∼ 54-56°C in the absence of DCMU, and ∼ 24°C in the presence of DCMU in PsbA1-cells (Panel E) and PsbA3-cells (Panel F), as already observed [38]. In PsbA1/S264V-cells (Panel G) the two TL curves were hardly affected when compared to PsbA1-cells. In contrast, in PsbA3/S264V-cells the TL was significantly modified when compared to PsbA3-cells with a peak at ∼ 44°C in the absence of DCMU instead of ∼ 54-56°C in PsbA3-cells, and at ∼ 38°C in the presence of DCMU instead of ∼ 24°C in PsbA3-cells. The peak observed at ∼ 68°C in the cells, more clearly seen in the presence of DCMU, likely originates from charge recombination between TyrD^●^ and QA^-^.

**Table 2.** Production of singlet oxygen with and without herbicides

| Strain | +DCMU |  | no DCMU |  | effect of DCMU |
| --- | --- | --- | --- | --- | --- |
| | $\mu\text{mol } ^1\text{O}_2 (\text{mg Chl})^{-1} \text{h}^{-1}$ | (%) | $\mu\text{mol } ^1\text{O}_2 (\text{mg Chl})^{-1} \text{h}^{-1}$ | (%) | % |
| WT*1 | 243 ± 5.7 | (100) | 287 ± 11 | (100) | 15 |
| PsbA1/S264V | 225 ± 6.8 | ( 92) | 265 ± 9.1 | ( 92) | 15 |
| WT*3 | 118 ± 5.7 | (100) | 144 ± 11 | (100) | 18 |
| PsbA3/S264V | 210 ± 8.5 | (179) | 211 ± 11 | (147) | 0 |

The TL curves after a single flash in the absence of DCMU (black curves) are expected to correspond to S2QB^-^ charge recombination. In PsbA1-PSII, PsbA3-PSII, and PsbA1/S264V-PSII, the TL was more or less as expected. The most striking observation is in PsbA3/S264V-PSII (Panel D), where the TL curve, with and without DCMU, exhibits characteristics close to those of the S2QA^-^/DCMU charge recombination in PsbA3-PSII (Panel B). The simplest explanation for this result is an inefficient electron transfer between QA^-^ and QB in the PsbA3/S264V-PSII mutant. This could be due to *i*) a more negative *E* (QB/QB^-^) or *ii*) the lack of QB. These hypothesis would align with EPR data, which show no Fe^2+^QB^-^ signal after a single flash in PsbA3/S264V-PSII. The differences between PsbA3/S264V-PSII and PsbA3/S264V-cells could be due to the large PQ pool in cells.

After 2 flashes (blue curves in Fig. 5), in PsbA1-PSII, PsbA1/S264V-PSII, and PsbA3-PSII, the TL curve corresponded well to a S3QB^-^ charge recombination, with a lower peak temperature than that for the S2QB^-^ charge recombination and a larger amplitude, as usually observed [52,53,56]. In PsbA3/S264V-PSII, the amplitude of the curve after two flashes was also larger than after one flash, but the peak temperature remained unchanged. This result is consistent with the understanding that, after the first flash, the electron is primarily used to reduce the oxidized non-heme iron, as demonstrated by EPR data and, after the second flash the TL corresponded *i*) to a S2QA^-^ charge recombination in centers with no oxidized non-heme iron prior to the flash, and *ii*) to a S3QA^-^ charge recombination in centers with the oxidized non-heme iron reduced by QA^-^ after the first flash and QA^-^ formed again on the second flash.

An additional challenge in interpreting the TL data arises from the potential effect of DCMU binding [57], which may differently modify the *E*_m_ of the QA/QA^-^ couple in the mutants compared to the wild type. This is particularly evident in PsbA3/S264V-PSII, where, as noted earlier, the herbicide was less effective, possibly due to differences in binding affinity. This is not surprising since, as mentioned in the Introduction, mutations of D1/S264 are known to affect the binding/efficiency of herbicides. Therefore, the efficiency of four different herbicides has been tested in the PsbA1/S264V-PSII and PsbA3/S264V-PSII.

Table 1 shows the oxygen-evolving activity either in the presence of DCBQ alone, or in the presence of DCBQ and the herbicides DCMU, Bromoxynil, Bromacil, or Atrazine (see Fig. S2 for the formulae of these compounds). Addition of DCMU inhibited O2-evolving activity to a similar extent in both PsbA1-PSII and PsbA1/S264V-PSII (92% and 91%, respectively). In PsbA3-PSII, the inhibition was also comparable (95%). The most pronounced difference was observed in PsbA3/S264V-PSII, where the activity was reduced by only 38%, compared to more than 90% inhibition in the other three cases. With the 3 other types of herbicide a similar observation could be done with a lower inhibition in the PsbA3/S264V-PSII than in PsbA3-PSII.

The changes in the energetics on the acceptor side in the mutants studied here could potentially impact the production of singlet oxygen [47]. Table 2 reports this production in purified PSII, both in the presence and absence of DCMU. The ^1^O2 production in PsbA1-PSII was twice that in PsbA3-PSII, both in the presence and absence of DCMU, as expected from the smaller energy gap between PheD1^-^QA and PheD1QA^-^ in PsbA3-PSII when compared to PsbA1-PSII [38,39]. Importantly, we have shown that the electron tunneling simulations of PSII electron transfers predicted a similar S2QA^-^/DCMU charge recombination in PsbA1-PSII and PsbA3-PSII [39] and a larger ^1^O2 production in PsbA1-PSII as it is experimentally observed. In PsbA1/S264V-PSII, singlet oxygen production was barely affected compared to PsbA1-PSII. In contrast, in PsbA3/S264V-PSII, ^1^O2 production significantly increased, both with and without DCMU. These data further demonstrate that the S264V mutation has more pronounced effects in PsbA3-PSII than in PsbA1-PSII (see the Discussion section for possible explanations of the results in Table 2).

From the effects of the S264V mutation in PsbA1 and PsbA3 described above, changes in the kinetics of QA^-^ reoxidation were also anticipated. To investigate these kinetics, we measured the amplitude of the PheD1 band shift, which is induced by the QA^-^ formation, as a function of time following flash illumination (Fig. 6). These measurements were conducted after the first flash applied to dark-adapted PSII, with PPBQ present to also ensure oxidation of the non-heme iron in a proportion of centers in the PsbA1/S264V-PSII mutant as shown in the EPR experiment (Fig. 4). We measured the amplitude of the 540 nm-*minus*-549 nm difference and the 543 nm-*minus*-553 nm difference in PSII containing either PsbA1 or PsbA3, respectively [39]. The spectra of these band shifts are shown in the supplementary material (Fig. S7). This measurement accounts for both the electron transfer from QA^-^ to Fe^3+^ as well as the electron transfer from QA^-^ to either QB or PPBQ.

**Figure 6:**
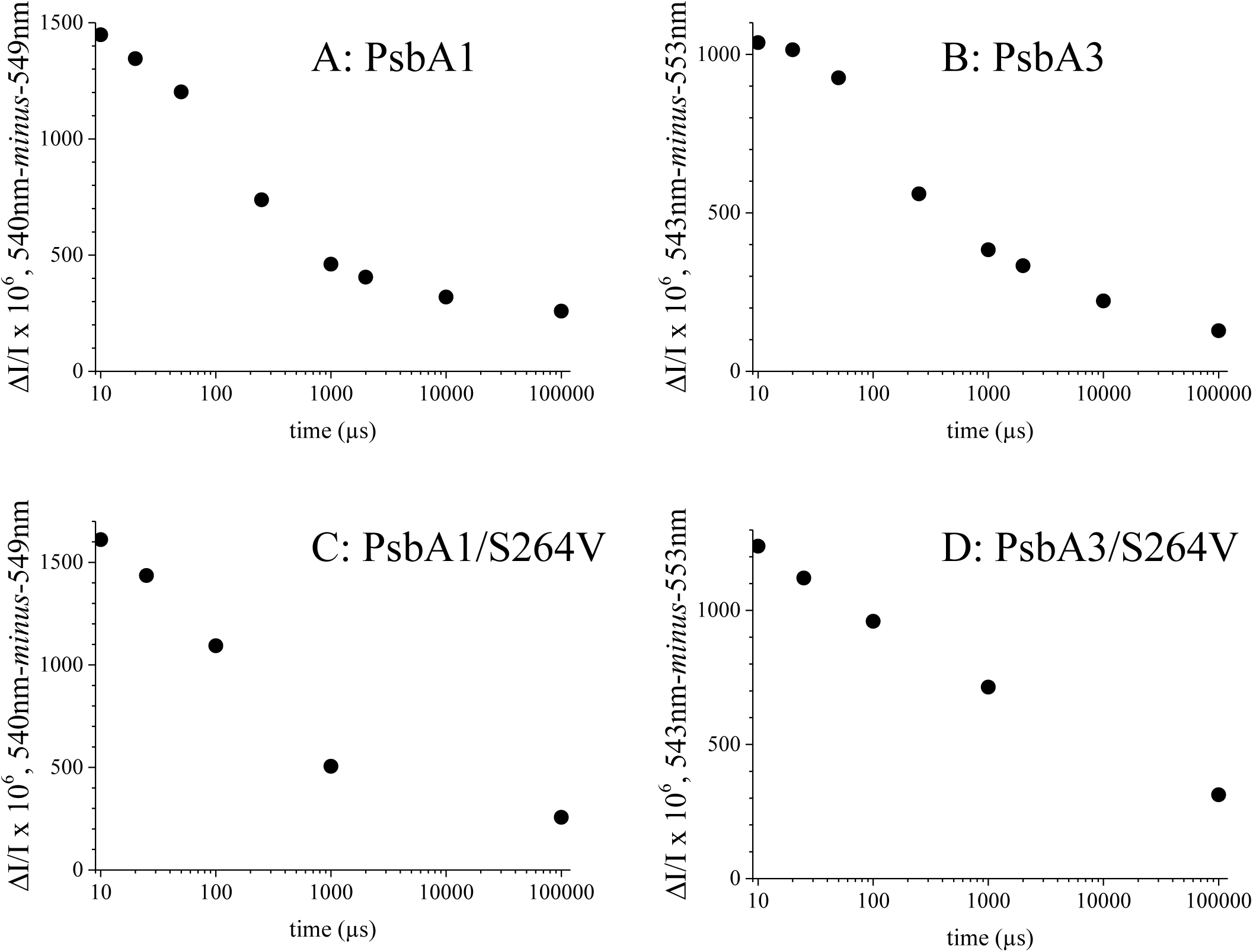
Time-courses of the absorption change differences either 540 nm-*minus*-549 nm in PsbA1-PSII (Panel A), and PsbA1/S264V-PSII (Panel C) or 543 nm-*minus*-553 nm in PsbA3-PSII (Panel B), and PsbA3/S264V-PSII (Panel D). 100 µM PPBQ dissolved in dimethyl sulfoxide were added to the samples after the dark-adaptation for 1 h at room temperature. The trace were normalized to a Chl concentration corresponding to *A*673 =1.75 (∼ 25 µg Chl mL^-1^).

In Panels A, B, and C, for the PsbA1-PSII, PsbA1/S264V-PSII, and PsbA3-PSII, respectively, the PheD1 band shift decay, and thus QA^-^ re-oxidation, exhibited global *t*1/2 values, between 300 µs and 400 µs. However, in the PsbA3/S264V-PSII, the global *t*1/2 was significantly longer than 1 ms, further highlighting its distinct behavior and reinforcing the hypothesis of a shift of the redox equilibria on the acceptor side in this mutant.

The electron transfer between QA^-^ and Fe^3+^ was estimated to be around 50 µs [37]. However, this measurement was conducted in the presence of DCMU, which is known to increase the midpoint potential of the QA/QA^-^ couple by approximately 50 mV in plant PSII [57], potentially slowing the rate of electron transfer between QA^-^ and Fe^3+^. It has been proposed that under certain conditions, this electron transfer between QA^-^ and Fe^3+^ can occur as rapidly as with a *t*1/2 ∼ 5-10 µs, as reviewed in [58]. In the experiment reported in Fig. 6, we also observed such a fast phase, between 2 µs and 10 µs, with comparable *t*1/2 and amplitudes across the four samples (these data are not shown in Fig. 6). However, as shown in Fig. S7, the spectrum of this fast phase (2 µs-*minus*-10 µs) was almost flat in the spectral range studied. This result strongly suggests that either QA^-^ was not involved or, within this time range, QA^-^ does not induce a spectral change of PheD1 but instead causes another spectral change with a more or less flat spectrum between 520 nm and 560 nm. In Fig. 6, nevertheless the limited number points, it seems that a fast phase is present for the PsbA3-PSII and PsbA1-PSII, with a decay compatible with a *t*1/2 of 50 µs. In the two mutants, considering the experimental accuracy, there is no evidence supporting the existence of such a fast phase.

The oxidation of non-heme iron is coupled to a proton release and it was proposed that D1/H215 is responsible for proton release during Fe^2+^ oxidation [33,34]. Conversely, it would be logical to assume that D1/H215 is also involved in the proton uptake during Fe^3+^ reduction. Therefore, we investigated the kinetics of proton uptake coupled to non-heme iron reduction in the two S264V mutants. For that, we followed the time-resolved absorption changes of bromocresol purple at 575 nm [59] after the first flash given to PsbA3-PSII (black), PsbA1/S264V-PSII (magenta), and PsbA3/S264V-PSII (blue), in the presence of PPBQ (Fig. 7).

**Figure 7:**
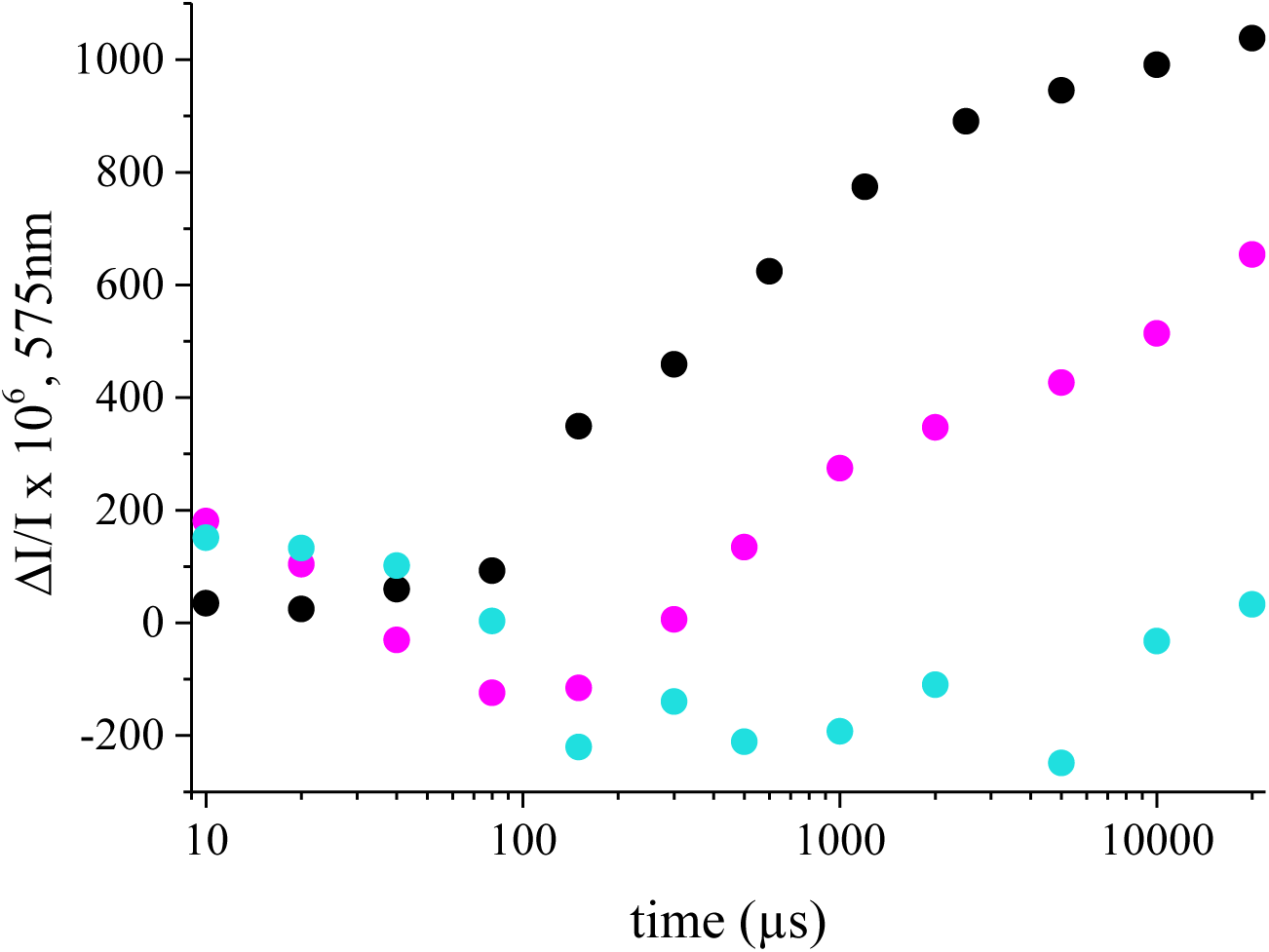
Time-courses of the absorption changes of bromocresol purple at 575 nm after the 1^st^ flash given to dark-adapted PsbA3-PSII (black points), PsbA1/S264V-PSII (red points), PsbA3/S264V-PSII (blue points). 100 µM PPBQ dissolved in dimethyl sulfoxide were added to the samples after the dark-adaptation for 1 h at room temperature.

Fig. 7 shows that proton uptake in PsbA3-PSII (black points) occurred with a *t*1/2 of approximately 400 µs. This *t*1/2 is significantly longer than the value proposed for electron transfer between QA^-^ and Fe^3+^ (equal or faster than 50 µs as discussed above) that suggests that the proton uptake coupled to this electron transfer occurs much after. In the PsbA1/S264V-PSII (magenta points), the proton uptake occurred with kinetics and amplitude similar to those in PsbA3-PSII, accounting for the baseline drift observed between 10 µs and 100 µs. If this drift has any significance, it may correspond to a proton release in a small proportion of centers, though this phenomenon has not been further investigated. In contrast, in PsbA3/S264V-PSII (blue points) no proton uptake was observed until 10 ms. The measurement was not extended beyond 20 ms due to the increasing dominance of baseline drifts at longer times. Therefore, it cannot be ruled out that proton uptake occurs after 20 ms in PsbA3/S264V-PSII. Nonetheless, this sample exhibits a distinctly marked phenotype compared to both PsbA3-PSII and PsbA1/S264V-PSII.

The potential absence of proton uptake (or strongly delayed) in PsbA3/S264V-PSII upon reduction of the oxidized non-heme iron may stem from a change in the *p*K of one or more groups involved in this reduction. Such *p*K changes imply the reorganization of the H-bond network in the Fe^2+/3+^QA region. One way to investigate this reorganization is by measuring the electrochromic response to QA^-^ formation [60]. To this end, we measured the spectral changes induced by illumination at 77 K in the absence of artificial electron acceptors (Fig. 8).

**Figure 8:**
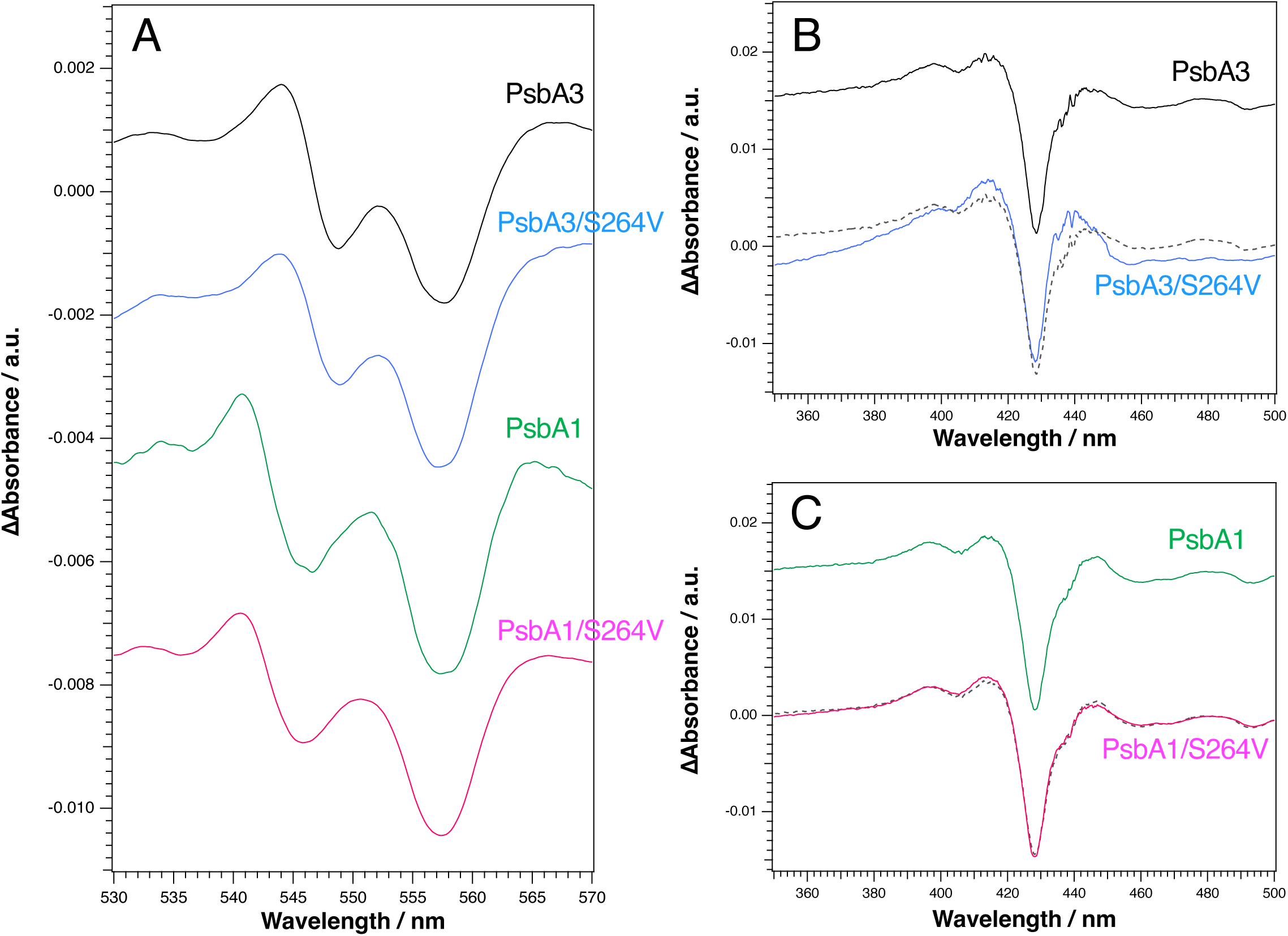
Light-*minus*-dark difference absorption spectra recorded at 77 K. After the recording of the spectrum on the dark-adapted samples, illumination at 77 K was given before the recording of the light spectrum on PsbA3-PSII (black spectra), PsbA3/S264V-PSII (blue spectra), PsbA1-PSII (green spectra), PsbA1/S264V-PSII (red spectra). Panel A shows the spectral region including the C-550 band-shift and the α-band of Cyt*b*559. Panels B and C show the spectral region including the reaction QY electrochromisms. The dashed lines in Panel B and C are a copy of the spectra in PsbA3-PSII and PsbA1-PSII, respectively.

Panel A in Fig. 8 shows to which extent the charge separation occurred at 77 K in the samples studied here by measuring the amplitude of the PheD1 band shift centered at 543 nm in PsbA1-PSII and PsbA1/S264V-PSII, and at 546 nm in PsbA3-PSII and PsbA3/S264V-PSII, as expected [39]. The amplitude of the band shift was comparable in PsbA1-PSII, PsbA1/S264V-PSII, and PsbA3-PSII, indicating a similar yield in the Q_A_^-^ formation. However, the band shift amplitude was slightly smaller in PsbA3/S264V-PSII, consistent with a proportion of PsbA3/S264V-PSII centers having an oxidized non-heme iron prior to the 77 K illumination. The presence of a negative peak at ∼557.5 nm in all samples, with a similar amplitude, confirms that Cyt*b*559 acted as the electron donor during 77 K illumination. The negative band at ∼ 557.5 nm arises from the smaller amplitude of the α-band absorption of Cyt*b* ^+^ compared to Cyt*b*559 [61].

Panel B in Fig. 8 presents the spectral changes in the Soret region induced by 77 K illumination in PsbA3-PSII (black spectrum) and PsbA3/S264V-PSII (blue spectrum). The large negative signal at ∼ 430 nm and the smaller positive signal at ∼ 415 nm correspond to changes in the Soret band of Cyt*b*559 upon its oxidation [61]. Between 440 nm and 460 nm - a region associated with electrochromic effects due to QA reduction - small differences between PsbA3-PSII and PsbA3/S264V-PSII were observed. These changes may reflect the reorganization of the hydrogen-bond network mentioned above. In contrast, Panel C shows that no differences were observed between PsbA1-PSII (green spectrum) and PsbA1/S264V-PSII (magenta spectrum).

## Discussion

To further investigate the differences between PSII with either PsbA1 or PsbA3, the D1/S264V mutants were constructed, with either D1 = PsbA1 or D1 = PsbA3. While the D1/S264V mutation with D1 = PsbA1 had far fewer consequences, although it was not entirely silent, the S264V mutation had significant consequences in PsbA3-PSII that can be summarized as it follows:

1. TL data supported the inefficient electron transfer from QA^-^ to QB;
2. kinetically, oxidation of QA^-^ was slowed, by at least an order of magnitude;
3. inhibition of O2 evolution activity by herbicides was weaker;
4. no Fe^2+^QB^-^ EPR signal was detected in dark-adapted PSII; instead,
5. a large Fe^3+^ EPR signal was present with
6. modifications in the shape of the Fe^3+^ EPR signal;
7. no QA^-^Fe^2+^QB^-^ biradical EPR signal was observed after a further illumination at 198 K following a flash illumination, confirming the lack of QB^-^ formation;
8. either no proton uptake coupled to non-heme iron reduction occurred or with a very slow rate compared to PsbA3-PSII;
9. changes were noted in the electrochromic response associated with Q_A_^-^ formation;
10. increased production of singlet oxygen, both with and without herbicides.

Given the known structures of PsbA3-PSII and PsbA1-PSII [2,62], the different consequences of the S264V mutation in these two PSII could merit further investigation using computational approaches in the future. It has been previously proposed that differences between PsbA1 and PsbA3 could arise, at least in part, from a long-distance effect involving the SQDG molecule [35]. In PsbA1-PSII, SQDG forms a hydrogen bond with D1/S270, whereas in PsbA3-PSII, the D1/A270 substitution prevents this interaction (Fig. 9) [2, 62].

**Figure 9:**
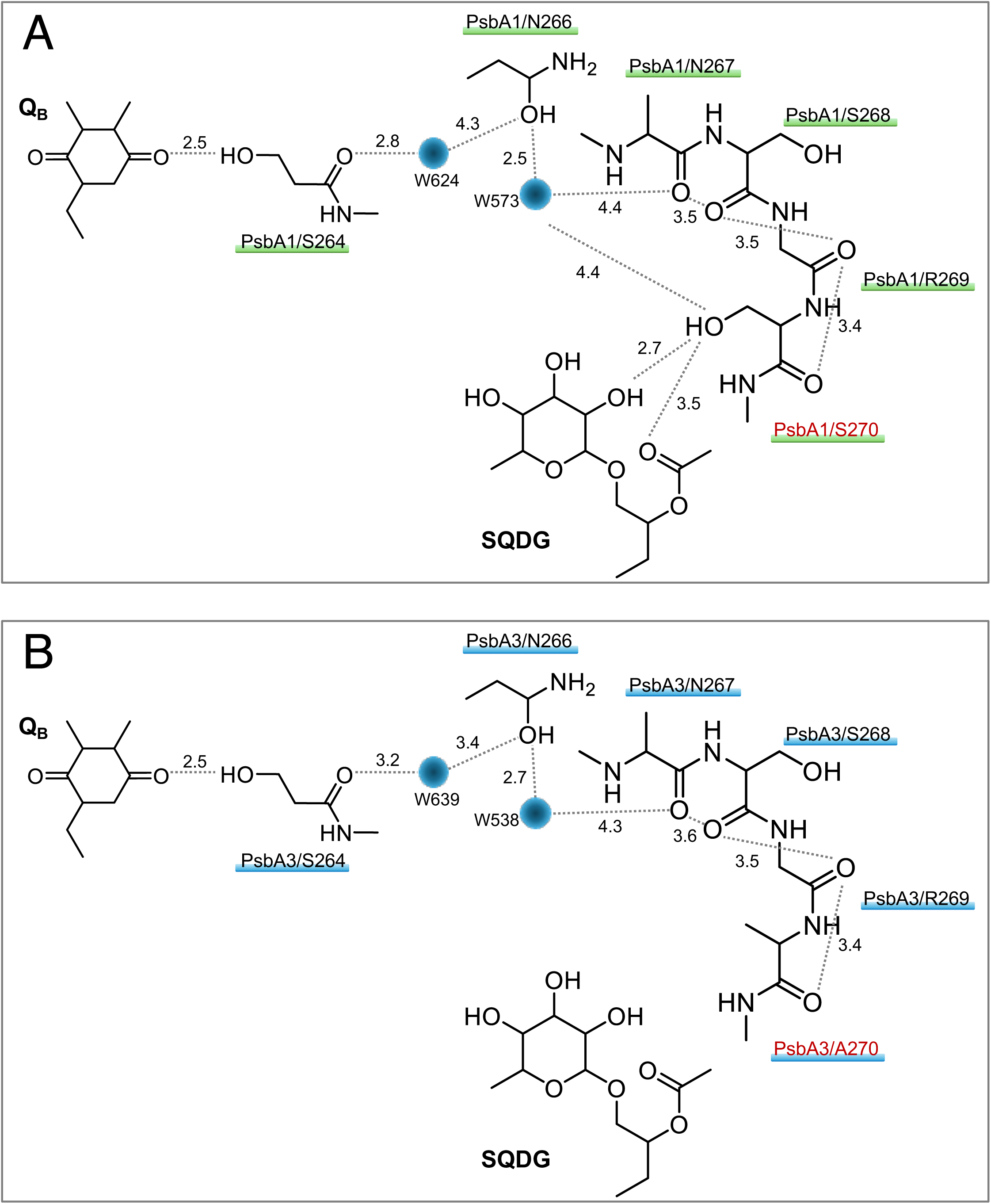
Comparison of hydrogen-bond network from QB to SQDG between PsbA1 (A) and PsbA3-PSII (B). The difference is that the amino acid at D1/270 is Ser in PsbA1 and Ala in PsbA3. PsbA1 and PsbA3. Therefore, in PsbA1, Ser270 makes hydrogen bonding to SQDG, but in PsbA3, no hydrogen-bond because of replaced by Ala. The figures were drawn with MacPyMOL with the A monomer of PsbA1-PSII (A) and PsbA3-PSII (B) in PDB 4UB6 [65] and PDB 7YQ7 [62], respectively.

This structural difference between PsbA1-PSII and PsbA3-PSII is thought to amplify the impact of the D1/S264V mutation.

Energetically, some differences has been found between PsbA1-PSII and PsbA3-PSII. In [15], the *E*_m_ of the QA^-^/QA couple in *T. elongatus* (in TyrD-less PSII *i.e.* with D1 = PsbA1) has been measured equal to -144 mV, that of the QB^-^/QB couple equal to +90 mV, that of the QBH2/QB^-^ couple equal to +40 mV, and that of the PQ pool equal to +117 mV. In other words, with a different experimental approach (spectroelectrochemistry *vs* EPR), the *E*_m_ of the QA^-^/QA couple in PsbA1-PSII from *T. elongatus* was found to be -140 mV [63], a value similar to that found in [15], and -89 mV in PsbA3-PSII [39]. As discussed in [15], the driving force for the electron transfer between QA^-^ and QB should be high enough to render this transfer efficient. We don’t know the *E*_m_ of the QB^-^/QB and QBH2/QB^-^ couples neither in PsbA3-PSII nor in PsbA3/S264V-PSII. However, the observation that the growth rate of PsbA3/S264V-cells was similar to that of PsbA3-cells is likely attributable to the presence of the large PQ pool in whole cells, which is absent in purified PSII in which 1 to 2 QB molecules are generally found [20]. This large PQ pool in cells likely helps maintain a sufficiently large driving force.

The redox characteristics of the acceptor side can be studied by TL [53–55]. In triazine-resistant mutants, as in *Erigeron canadensis* L, the TL curves after one flash exhibit a similar peak temperature with and without DCMU [64] as observed here in the PsbA3/S264V-PSII. In these triazine-resistant mutants, there is a mutation of the D1/S264 residue [29]. In [64], it was proposed that the D1/S264G mutation induced a strong shift to the left of the equilibrium QA^-^QB ⇌ QAQB^-^ due to a more negative potential for the QB/QB^-^ couple. This change in the QA^-^QB ⇌ QAQB^-^ equilibrium corresponds to a decrease in the energy gap between QA^-^QB and QAQB^-^ (see Fig. S10 for a simplified energetic scheme). All other things being equal, and applying the same reasoning as in [39], the increased production of ^1^O2 in PsbA3/S264V-PSII suggests a reduction in the energy gap between PheD1QA^-^ and PheD1^-^QA^-^ compared to PsbA3-PSII. This is at odds with the TL data which would suggest a small increase in the energy gap between PheD1^-^QA/DCMU and PheD1QA^-^/DCMU. However, we have seen in [38,39] that we must take all the recombination routes for making this kind of correlation.

The reduction of QB has been proposed to be coupled with the protonation of D1/H252 [33,34], and to a reorientation of the hydroxyl group of D1/S264 toward the distal QB carbonyl group. This structural change would stabilize QB^-^ and thereby facilitate electron transfer from QA^-^ to QB [16]. Our data provide experimental support for this model. In the PsbA3/S264V mutant, the distal QB carbonyl group cannot interact with a hydroxyl group, resulting in QB^-^ no longer being stabilized. Consequently, electron transfer from QA^-^ to QB is significantly slowed. This slowing down would correspond to several of the observations made here in the PsbA3/S264V-PSII.

A comparison of the X-ray crystal structures of PsbA1-PSII (*T. vulcanus*, PDB: 4UB6 [65]) and PsbA3-PSII (*T. elongatus*, PDB: 7YQ7 [62]) reveals the presence of a water molecule (W501) near bicarbonate and D1/Y246 in PsbA3-PSII, which is absent in PsbA1-PSII (Fig. S8). W501 is within hydrogen-bonding distance to both D1/Y246 and bicarbonate, suggesting a slightly enhanced proton transport pathway in PsbA3-PSII compared to PsbA1-PSII. Although W501 is located near D1/H215 and QB, it is too distant to form a direct hydrogen bond. The absence of electron density corresponding to W501 in the PsbA1-PSII crystal structure may not be due to species differences between *T. elongatus* and *T. vulcanus*, but rather to differences due to the PsbA1/PsbA3 exchange. Our findings suggest that the S264V mutation in PsbA3-PSII of *T. elongatus* disrupts the binding of non-heme iron and bicarbonate, implying that W501 helps maintain the structural integrity around QB, at least in PsbA3. In contrast, the same mutation in PsbA1-PSII did not cause such severe effects, suggesting that PsbA1 may lack a water molecule equivalent to W501, similar to PSII in *T. vulcanus*.

The cryo-EM structure of PsbA1-PSII in *T. elongatus* reported by Hussein *et al.* [66], reveals a water molecule, W538 (referred to as W63 in their paper), corresponding to W501 (Fig. S9, Panel C). However, the distance between this water molecule and bicarbonate (3.9 Å) is longer than that observed in the PsbA3-PSII structure reported by Shen *et al.* (3.2 Å), making hydrogen bonding less likely. This suggests that even if PsbA1-PSII contains W501 (W538), its interaction with QB may be weak. Furthermore, as shown in Fig. S9 and Table S1, the presence or absence of water molecules in the electron acceptor side not only differs between PsbA1- and PsbA3-PSII but also varies between the A and B monomers within the same PSII complexes. These differences may be attributed to the structural flexibility of this region, which accommodates QB quinone entry and exit and is characterized by high B-factor values. Variations in water molecule positioning among purified samples suggest that hydrogen bonding between PQ and water molecules may be disrupted during PQ movement *in vivo*. To prevent inhibition of proton transfer under such conditions, the hydrogen-bonding network in this region may have evolved to establish multiple proton transport pathways, allowing for detours around potential “blocked” routes. Further studies are needed to clarify the role of W501 in proton transport.

The reduced efficiency of DCMU addition seen in Table 1 is likely due to the modification of the QB binding site. Despite the high concentration of DCMU, binding may still occur in all centers, but with less effective inhibition of electron transfer. In other words, the binding mechanism may differ between PsbA3/S264V-PSII and PsbA3-PSII. For the other herbicides, the percentage of inhibition observed in PsbA1-PSII, PsbA1/S264V-PSII, and PsbA3-PSII was similar to that seen with DCMU. However, in PsbA3/S264V-PSII, the inhibition was slightly greater. We hypothesize that this result arises because the binding sites for these herbicides do not perfectly match the QB binding in its site, making the structural modifications in the mutant less impactful on herbicide binding.

The significantly lower oxygen evolution activity of PsbA3/S264V-PSII compared to PsbA3-PSII in the presence of DCBQ (Table 1) suggests that DCBQ binding at the QB site, similar to the binding of QB itself [67], is altered in this mutant. A modification of the QB binding site is expected to affect the redox properties of QB, as discussed above likely destabilizing QB^-^ in PsbA3/S264V-PSII.

The efficiency of various types of herbicides, such as DCMU, bromacil, bromoxynil, and triazine, in inhibiting O2 evolution is clearly illustrated in Table 1 for PsbA1-PSII, PsbA3-PSII, and PsbA1/S264V-PSII. However, in PsbA3/S264V-PSII, the inhibition is reduced, particularly with DCMU, which inhibits only 38% of the O2 evolution. These results suggest that the S264V mutation in PsbA3, by disrupting the hydrogen bond with QB, alters the structure of the QB-binding pocket, thereby affecting the binding strength of herbicides with different structures.

In dark-adapted PsbA1-PSII and PsbA3-PSII from *T. elongatus*, approximately 50% of the centers are in the Fe^2+^QB^-^ state, while the other 50% are in the Fe²⁺ QB state [20,37]. In PsbA1/S264V-PSII, the Fe^2+^QB^-^ EPR signal is smaller than in PsbA1-PSII, and in PsbA3/S264V-PSII it is not detected. However, the amplitude of the Fe^2+^QB^-^ EPR signal was not precisely quantified. Accurate quantitation would require a large amount of PSII and, for example, a period-two oscillation experiment with sequential flashes (*e.g.*, 1, 2, 3, 4), which was not performed here.

In plant PSII, QB^-^ is unable to oxidize the non-heme iron. However, PPBQ^-^ can do so [31]. In *T. elongatus*, QB^-^ is capable of oxidizing the non-heme iron, suggesting that QB^-^ can shift to a position close to its site that raises the *E* of the QB^-^/QBH2 couple from 40 mV to above that of the Fe^3+^/Fe^2+^ couple (∼ 400 mV, [68]). In PsbA3/S264V-PSII, either the *E* of the QB^-^/QBH2 couple in this alternative site is even higher, or the *E* of the Fe^3+^/Fe^2+^ couple is lower, which would account for the larger amplitude of the non-heme iron EPR signal and the absence of the Fe^2+^QB^-^ EPR signal. We hypothesize that upon the oxidation of Fe^2+^ to Fe^3+^, the QBH2 formed leaves its binding site. In whole PsbA3/S264V cells, QBH2 is replaced by an oxidized plastoquinone (PQ) from the pool, resulting in a TL curve that could be attributable to the S2QB^-^ charge recombination after one flash in the absence of DCMU. In contrast, in PsbA3/S264V-PSII, there is not enough oxidized PQ molecule available to replace the QBH2 that has left the site. This results in a situation where the non-heme iron remains in the oxidized state (Fe^3+^) without a bound QB.

The EPR in PsbA3/S264V-PSII revealed a structural heterogeneity at the Fe^3+^ site, with two structures corresponding to an Fe^3+^ reducible at different temperatures. Such temperature-dependent photoreduction heterogeneity has been reported previously in plant PSII [69]. The axial non-heme iron signal detected here in PsbA3/S264V-PSII is comparable to that observed in plant PSII in the presence of *o*-phenanthroline [51], where it was logically attributed to increased symmetry of the four histidine ligands possibly resulting in a change in the redox potential of the Fe^3+^/Fe^2+^ couple.

The data in Fig. 7 indicate that in PsbA3/S264V-PSII, there is either no proton uptake coupled to the reduction of the oxidized non-heme iron by QA^-^, or the proton uptake occurs much more slowly than in the other samples. One hypothesis could be that this lack of protonation is responsible for the different Fe^3+^ structure, which in turn accounts for the modified Fe^3+^ EPR signal. D1/H215 has been proposed as the residue responsible for proton release upon Fe^2+^ oxidation [33,34]. It is therefore plausible that D1/H215 also mediates proton uptake during Fe^3+^ reduction. Given that D1/H215 bridges the non-heme iron and QB (Fig. 1), any changes in the QB site, such as those induced by the S264V mutation in PsbA3-PSII, could impact the properties of D1/H215.

The *p*K for the deprotonation of D1/H215 in the Fe^3+^ state has been estimated to be ∼5.5 [34]. Since proton uptake in the Fe^2+^ state is observed at pH 6.5 in wild-type PSII, the *pK* for D1/H215 protonation in the Fe^2+^ state is expected to be > 6.5. In the PsbA3/S264V-PSII mutant, the *p*K of D1/H215 in the Fe^2+^ state may be even higher than in PsbA3-PSII. Additionally, it was proposed in [34] that the NπH proton of D1/H215 is highly sensitive to its electrostatic environment. Fig. 8 indeed suggests a change in the electrostatic environment of FeQA in the PsbA3/S264V-PSII mutant, supporting this hypothesis.

## Material

Additional EPR data, kinetics and structural analysis are shown.

## Acknowledgements

We thank to Yuki Ito (Ehime University) for technical assistance. MS was supported by a JSPS-KAKENHI grant 21H02447 and 24K21853. AB was supported in part by the French Infrastructure for Integrated Structural Biology (FRISBI) ANR-10-INBS-05. JS was supported by the Labex Dynamo (ANR-11-LABX-0011-01).

## Abbreviations

Photosystem II: PSII
Chl: chlorophyll
ChlD1/ChlD2: monomeric Chl on the D1 or D2 side, respectively
P680: primary electron donor
PD1 and PD2: individual Chl on the D1 or D2 side, respectively, which constitute a pair of Chl with partially overlapping aromatic rings
PheD1 and PheD2: pheophytin on the D1 or D2 side, respectively
QA: primary quinone acceptor
QB: secondary quinone acceptor
PQ: plastoquinone
TyrZ: the tyrosine 161 of D1 acting as the electron donor to P680
MES: 2-(*N*-morpholino) ethanesulfonic acid
PPBQ: phenyl *p*–benzoquinone
DCBQ: 2,6-dichloro-*p*-benzoquinone
DCMU: 3-(3,4-dichlorophenyl)-1, 1-dimethylurea
Bromoxynil: 3,5-dibromo-4-hydroxybenzonitrile
Bromacil: 5-bromo-6-methyl-3-(1-methylpropyl)-2,4(1H,3H)-pyrimidinedione
Atrazine: 6-Chloro-*N*-ethyl-*N’*-(1-methylethyl)-1,3,5-triazine
TL: thermoluminescence *Thermosynechococcus elongatus T. elongatus*
WT*3: *T. elongatus* mutant strain deleted *psbA1* and *psbA2* genes and with a His-tag on the carboxy terminus of CP43
WT*1: *T. elongatus* mutant strain deleted *psbA2* and *psbA3* genes and with a His-tag on the carboxy terminus of CP43
EPR: Electron Paramagnetic Resonance
PCET: proton coupled electron transfer
SQDG: sulfoquinovosyldiacylglycerol
*E*_m_: redox potential

---

**Figure S1.**
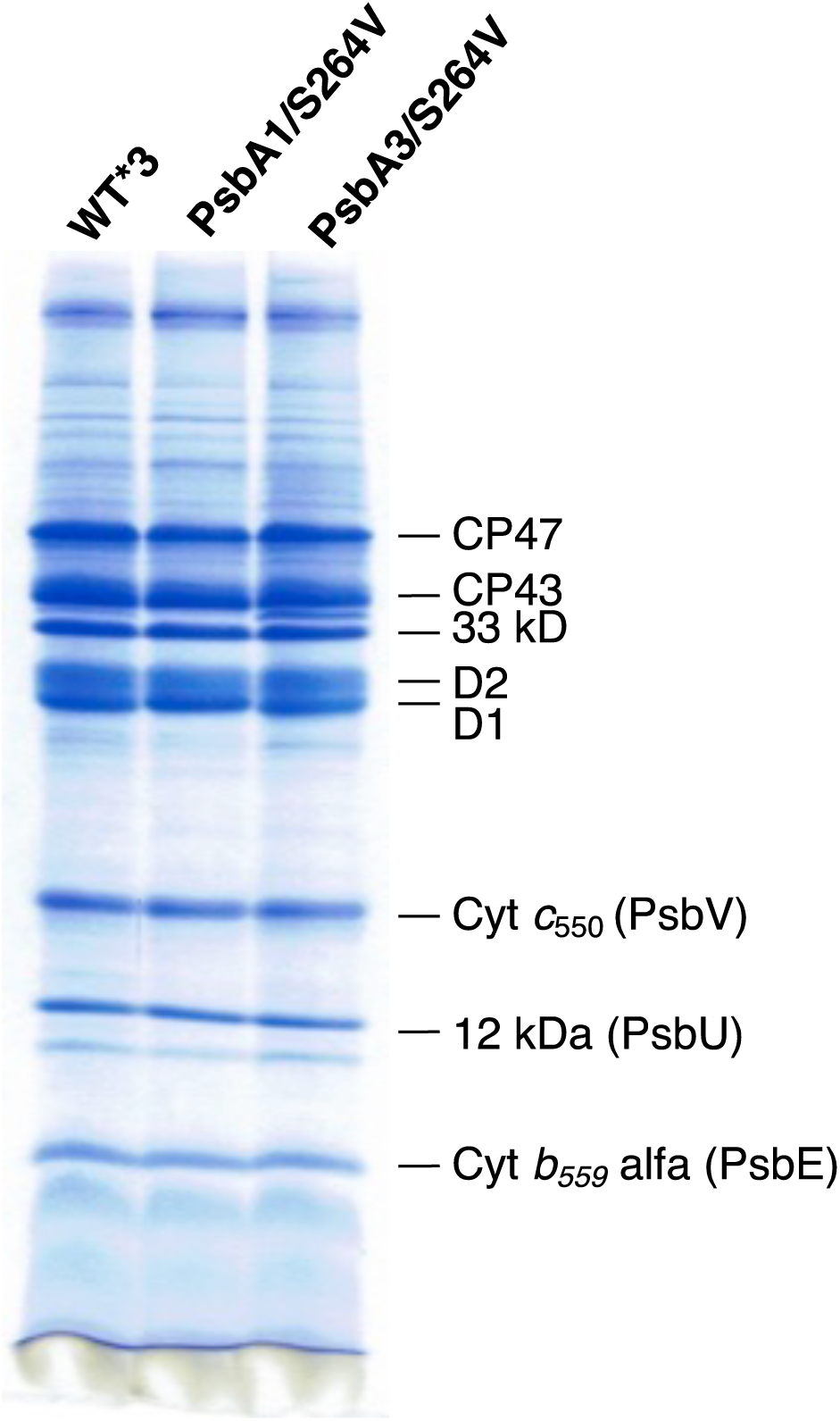
Coomassie brilliant blue-staining of SDS-polyacrylamide gel electrophoresis of PsbA3-PSII (WT*3), PsbA1/S264V-PSII, and PsbA3/S264V-PSII. The amount of PSII loaded was 8 μg of Chl for each lane.

**Figure S2.**
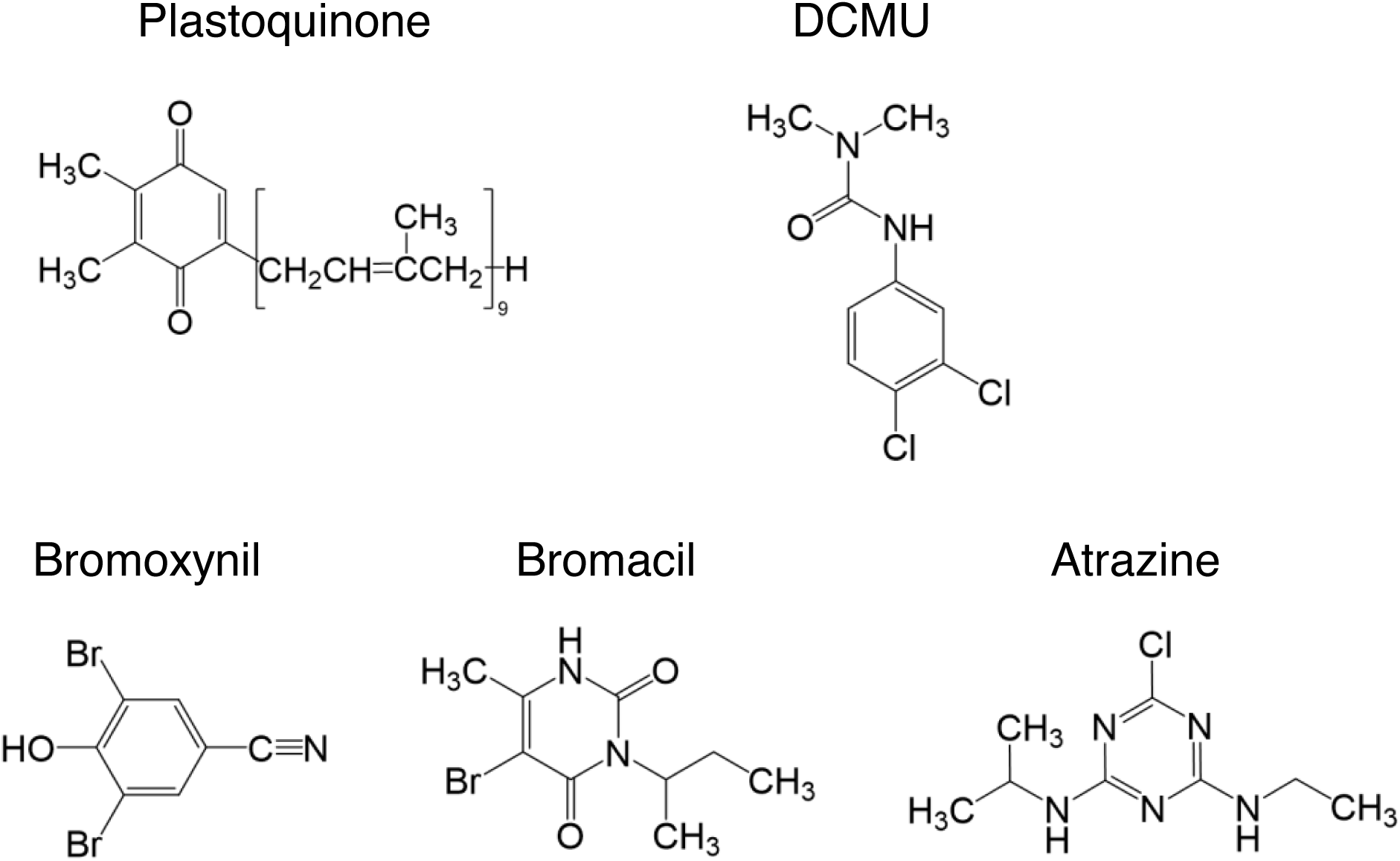
Skeletal formula of PQ9 (QA and QB), and of the herbicides DCMU, Atrazine, Bromoxynil, and Bromacil.

**Figure S3.**
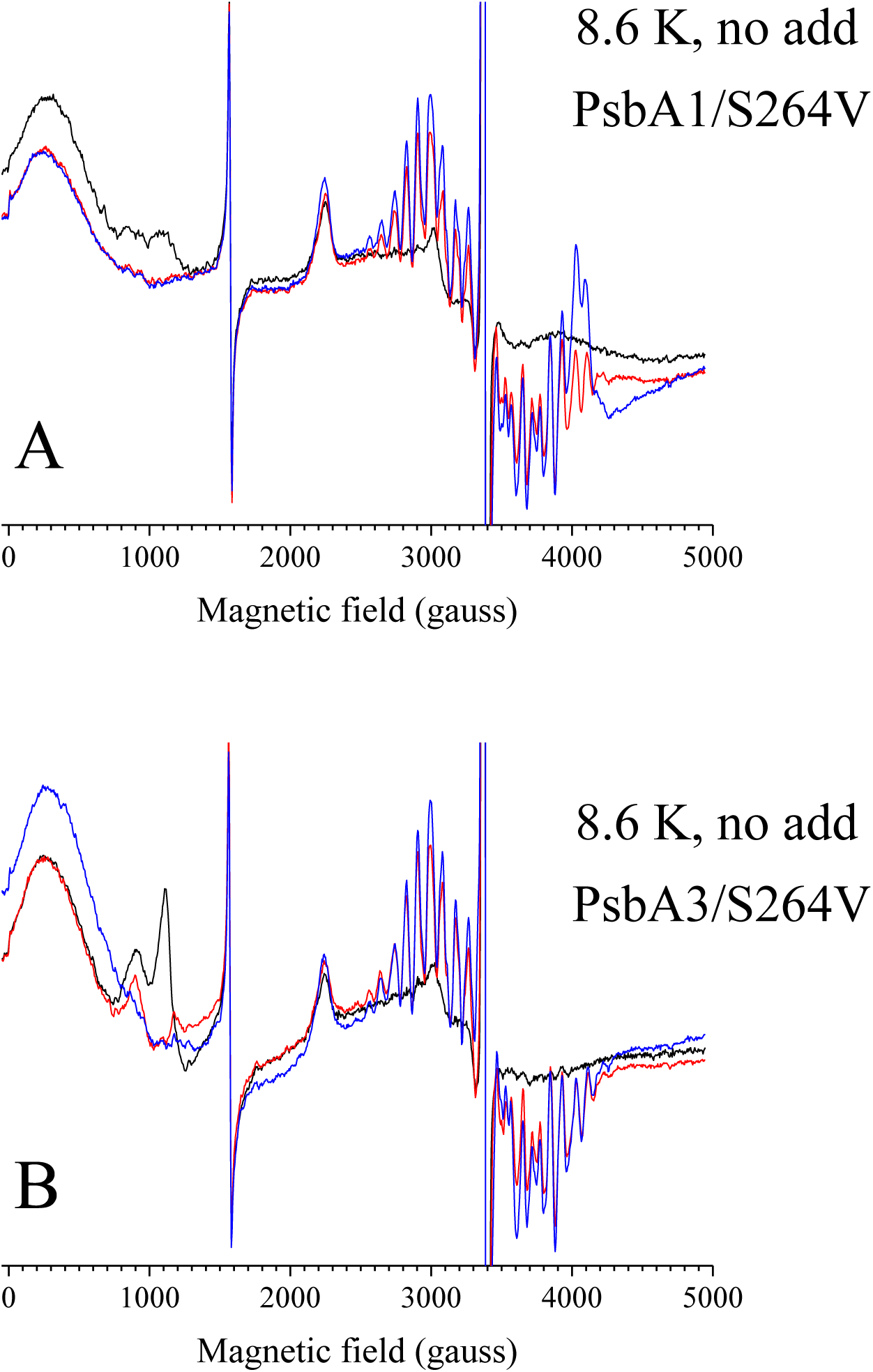
cw-EPR spectra recorded at 8.6 K in active PsbA1/S264V-PSII (Panel A) and active PsbA3/S264V-PSII (Panel B). Spectra in black were recorded in the dark-adapted samples. Spectra in red were recorded after laser flash illumination. The blue spectra were recorded after a further continuous white light illumination for 5-10 s at 198 K. The microwave power was 20 mW. The modulation frequency was 100 kHz, the modulation amplitude was 25 gauss, and the microwave frequency was 9.49 GHz.

**Figure S4.**
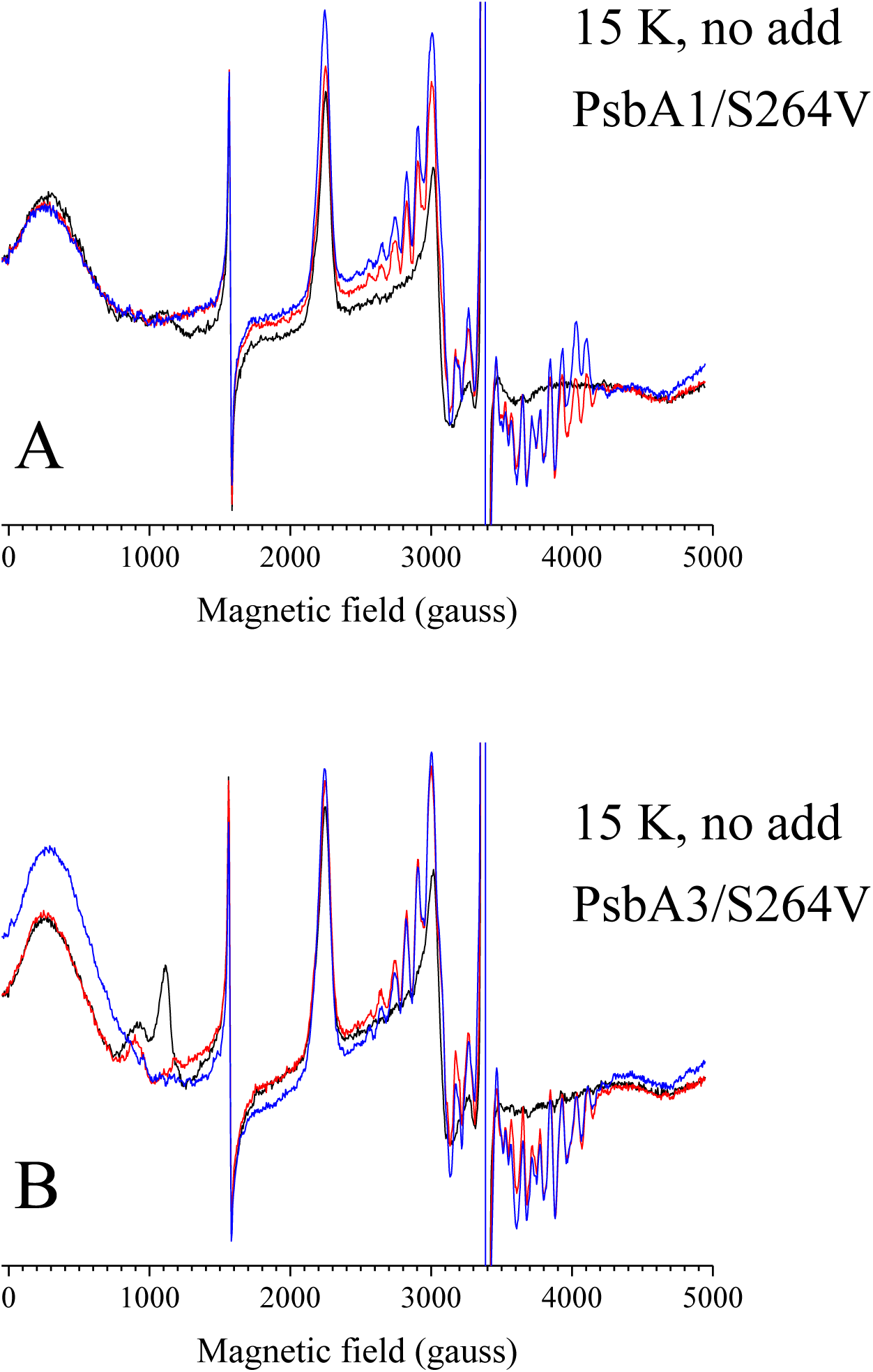
cw-EPR spectra recorded at 15 K in active PsbA1/S264V-PSII (Panel A) and active PsbA3/S264V-PSII (Panel B). Spectra in black were recorded in the dark-adapted samples. Spectra in red were recorded after laser flash illumination. The blue spectra were recorded after a further continuous white light illumination for 5-10 s at 198 K. The microwave power was 5 mW. The modulation frequency was 100 kHz, the modulation amplitude was 25 gauss, and the microwave frequency was 9.49 GHz.

**Figure S5.**
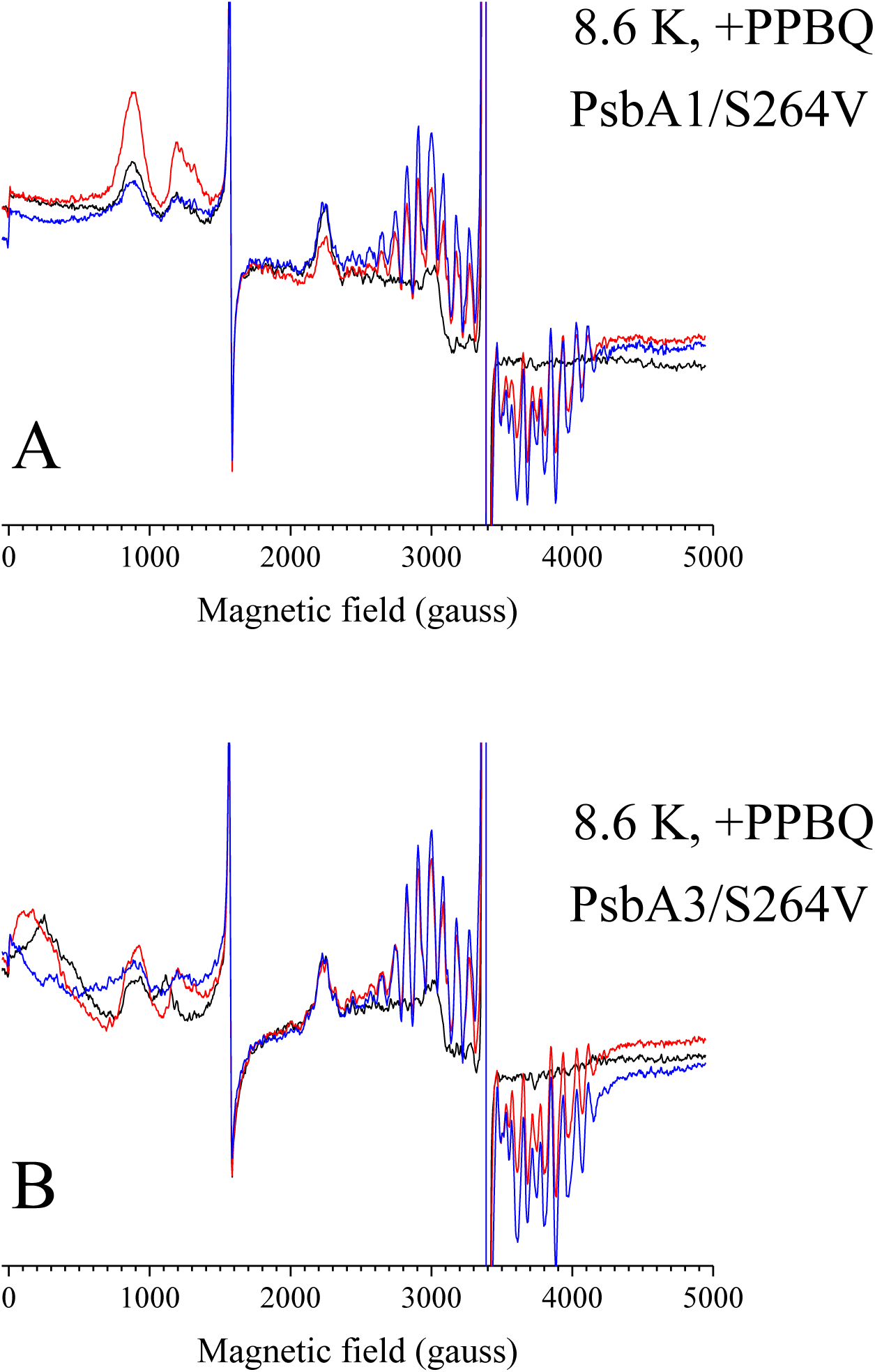
cw-EPR spectra recorded at 8.6 K in active PsbA1/S264V-PSII (Panel A) and active PsbA3/S264V-PSII (Panel B) in the presence of PPBQ. Spectra in black were recorded in the dark-adapted samples. Spectra in red were recorded after laser flash illumination. The blue spectra were recorded after a further continuous white light illumination for 5-10 s at 198 K. The microwave power was 20 mW. The modulation frequency was 100 kHz, the modulation amplitude was 25 gauss, and the microwave frequency was 9.49 GHz.

**Figure S6.**
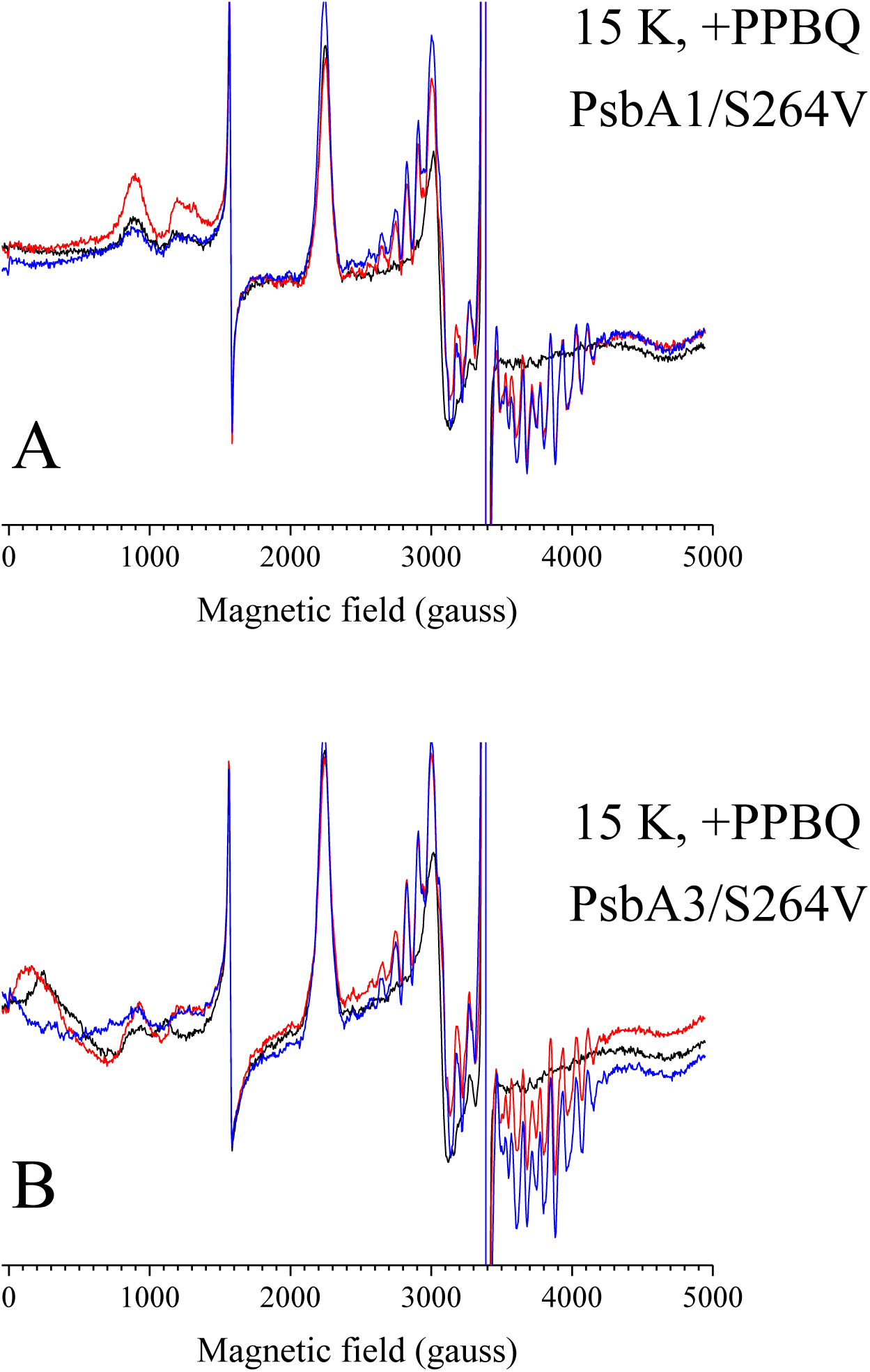
cw-EPR spectra recorded at 15 K in active PsbA1/S264V-PSII (Panel A) and active PsbA3/S264V-PSII (Panel B) in the presence of PPBQ. Spectra in black were recorded in the dark-adapted samples. Spectra in red were recorded after laser flash illumination. The blue spectra were recorded after a further continuous white light illumination for 5-10 s at 198 K. The microwave power was 5 mW. The modulation frequency was 100 kHz, the modulation amplitude was 25 gauss, and the microwave frequency was 9.49 GHz.

## Additional comments on Fig. S3 to S6

Spectra recorded at 8.6 K after a flash illumination at room temperature, both with and without PPBQ, exhibit an S2 multiline signal in both mutants with similar amplitude.

Additional continuous illumination at 198 K resulted in a slight increase in the multiline signal, consistent with a miss parameter of approximately 11%.

In the absence of PPBQ, as observed at 4.2 K, the oxidized non-heme iron signal recorded in the dark-adapted samples (black spectra) shows a much larger amplitude in PsbA3/S264V-PSII compared to PsbA1/S264V-PSII.

In spectra recorded at 15 K from dark-adapted samples, the resonances at ∼2250 gauss, ∼3075 gauss, and ∼4677 gauss correspond to the *g*z (∼3.02), *g*y (∼2.21), and *g*x (∼1.45) of Cyt*c*550. In the difference spectrum (not shown) corresponding to “1 flash + hν198K”-*minus*-“1 flash,” a small cytochrome signal was induced. The resonances at ∼2230 gauss, ∼3114 gauss, and ∼4725 gauss correspond to the *g*z (∼3.04), *g*y (∼2.18), and *g*x (∼1.44) of Cyt*b*559, which is likely oxidized in some centers already in the S2 state before the illumination at 198 K.

**Figure S7.**
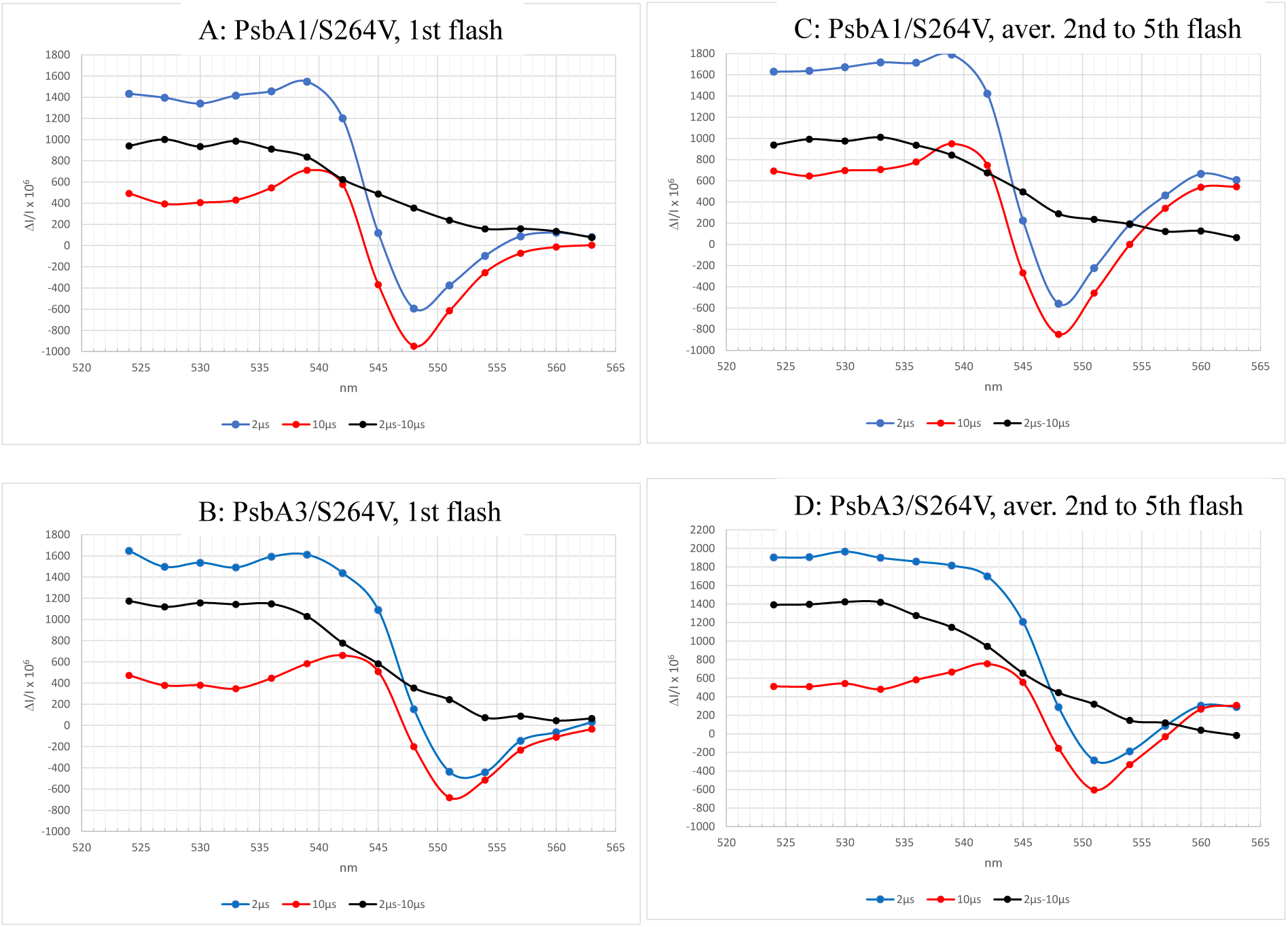
Light-induced difference spectra around 545 nm. The flash-induced absorption changes, in the presence of 100 µM PPBQ, were measured in PsbA1/S264V-PSII (Panels A and C), and PsbA3/S264V-PSII (Panels B and D). The spectra correspond either to the ΔI/I measured after the first flash (Panels A and B), or to the average of the spectra from the 2^nd^ to 5^th^ flash (Panels C and D).The Chl concentration was 25 μg mL^−1^. The blue difference spectra were measured 2 µs after the flash. The red spectra were measured 10 µs after the flash. The black spectra correspond to the difference 2 µs *minus* 10 µs. The red spectra well corresponds to the electrochromic blue shift undergone by PheoD1 upon reduction of QA and known as the C-550 bandshift. This bandshift is redshifted in PsbA3-PSII, and PsbA3/S264V-PSII when compared to PsbA1-PSII, and PsbA1-S264V, due to the glutamate at the position 130 in PsbA3 instead of a glutamine in PsbA1. In contrast, the signal which rapidly disappears between 2 µs and 10 µs is almost flat in this spectral region with no differences between PsbA1 and PsbA3. In this time scale, QA-is stable and the origin of this signal remains to be identified.

**Figure S8.**
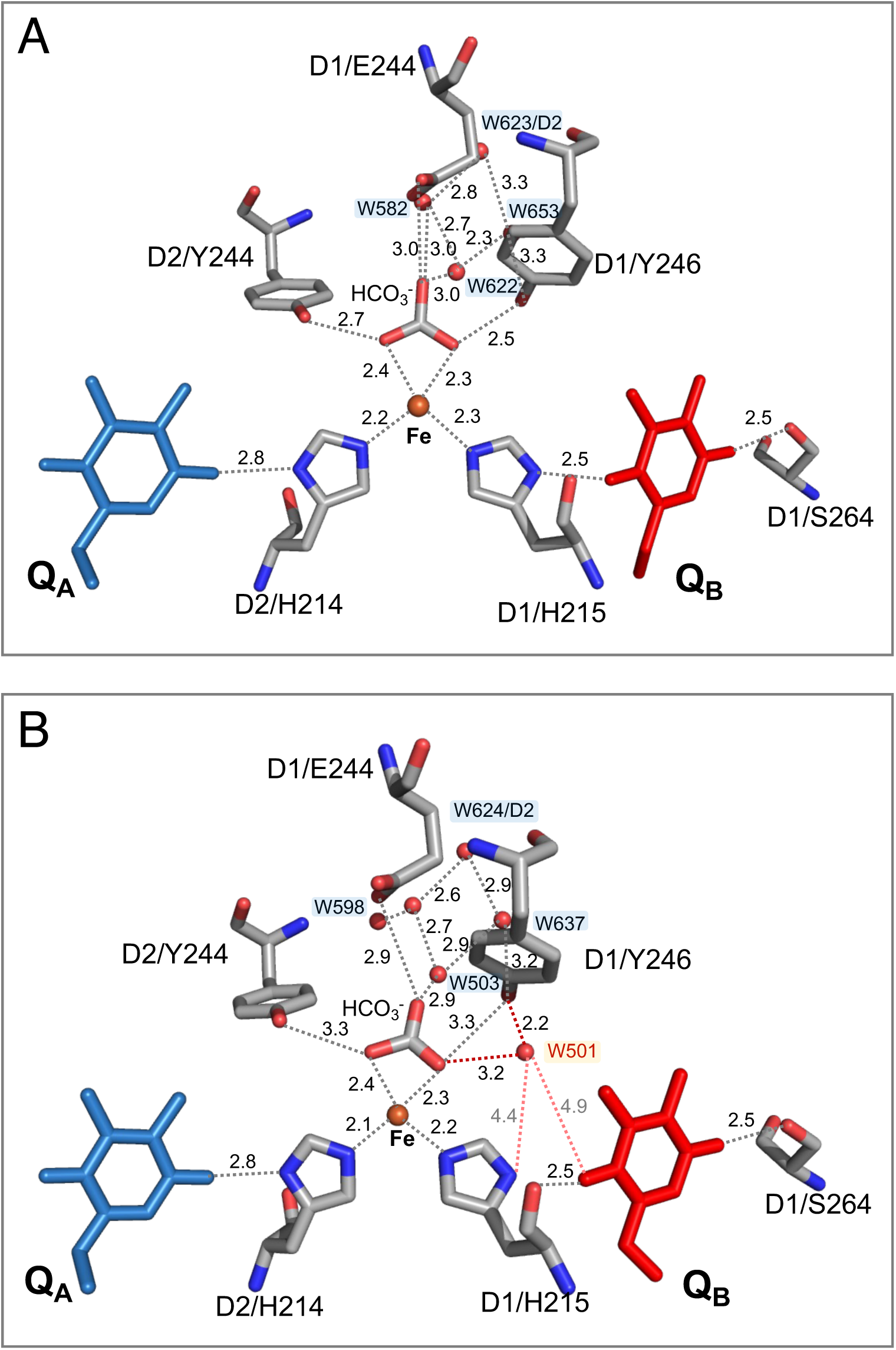
Comparison of hydrogen-bond network around acceptor side between PsbA1 (Panel A) and PsbA3-PSII (Panel B). In PsbA3, an extra water molecule, W501, exists near the bicarbonate and D1/Y246, forming hydrogen bonds with them, whereas there is no equivalent water molecule in PsbA1. The figures were drawn with MacPyMOL with the A monomer of PsbA1-PSII (A) and PsbA3-PSII (B) in PDB 4UB6 [61] and PDB 7YQ7 [62], respectively.

**Figure S9.**
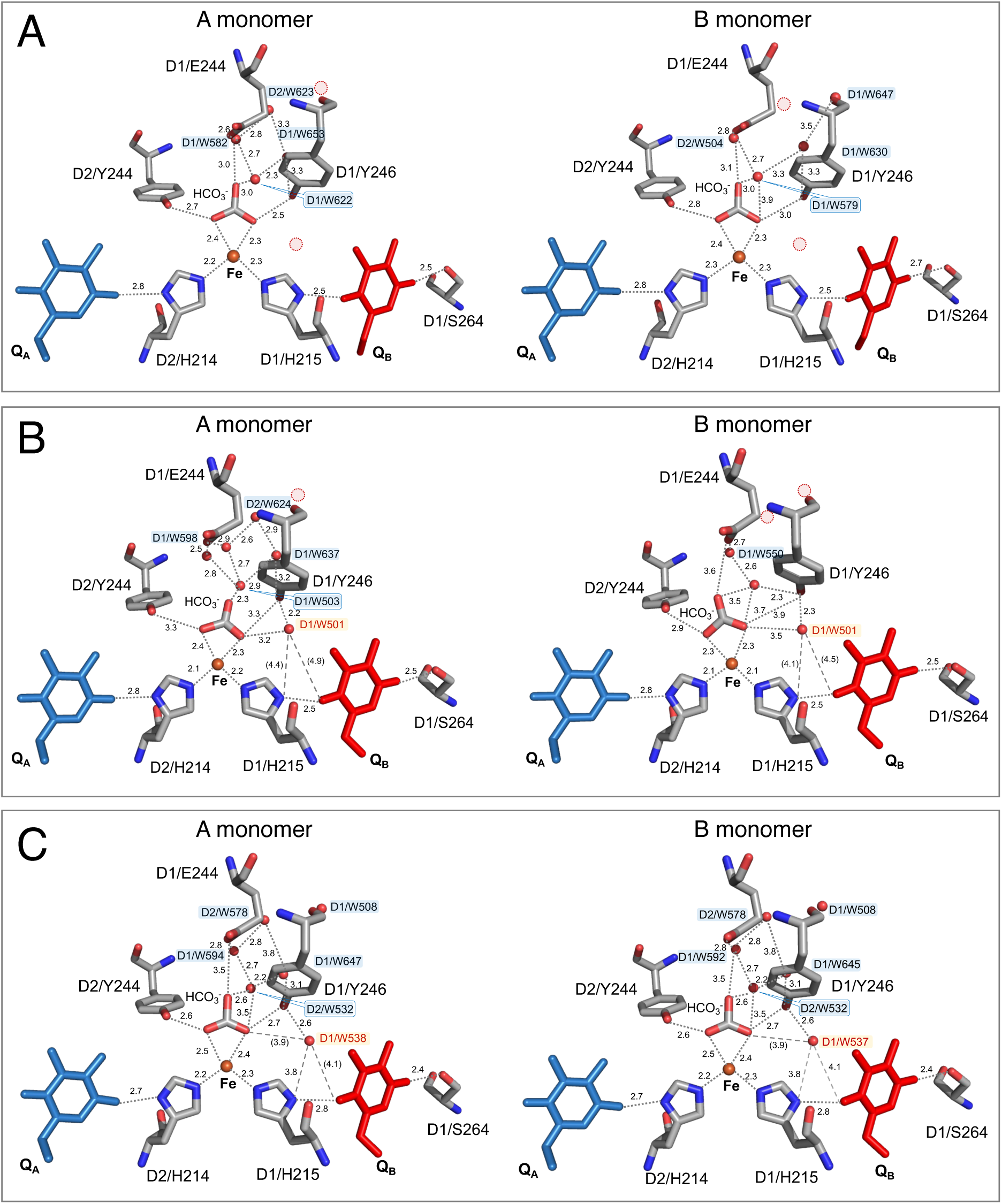
Comparison of water molecules and hydrogen-bond network around acceptor side among PDBs 4UB6 (PsbA1-PSII: Panel A) [61], 7YQ7 (PsbA3-PSII: Panel B) [62] and 9EVX (PsbA1-PSII: Panel C) [66]. Red circles indicates water molecules that are present in other structure.

**Figure S10.**
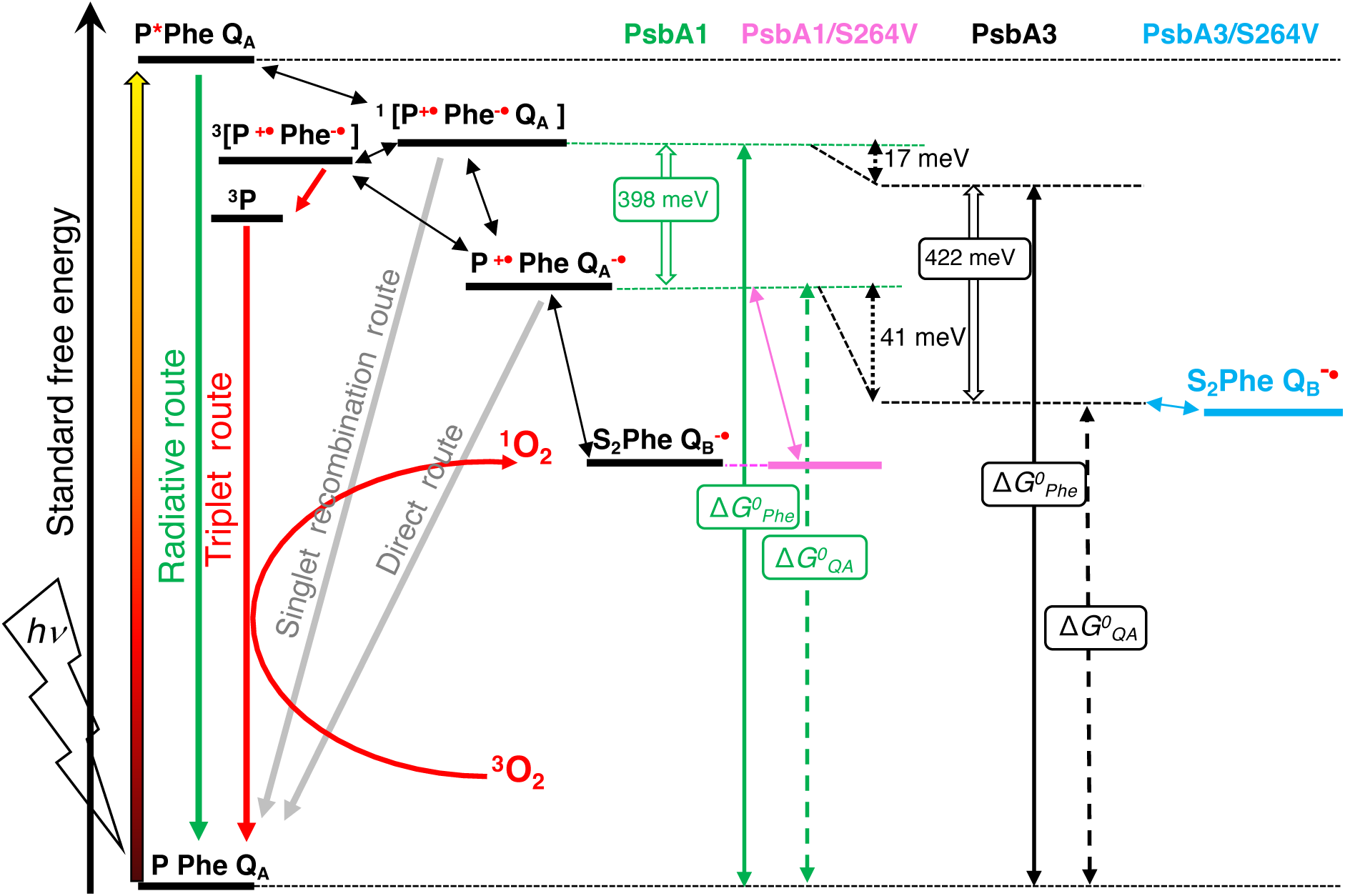
Simplified energetics scheme (from Ref. 39) showing the energy level of the QB/QB^-^ in the two mutants.

**Table S1.**
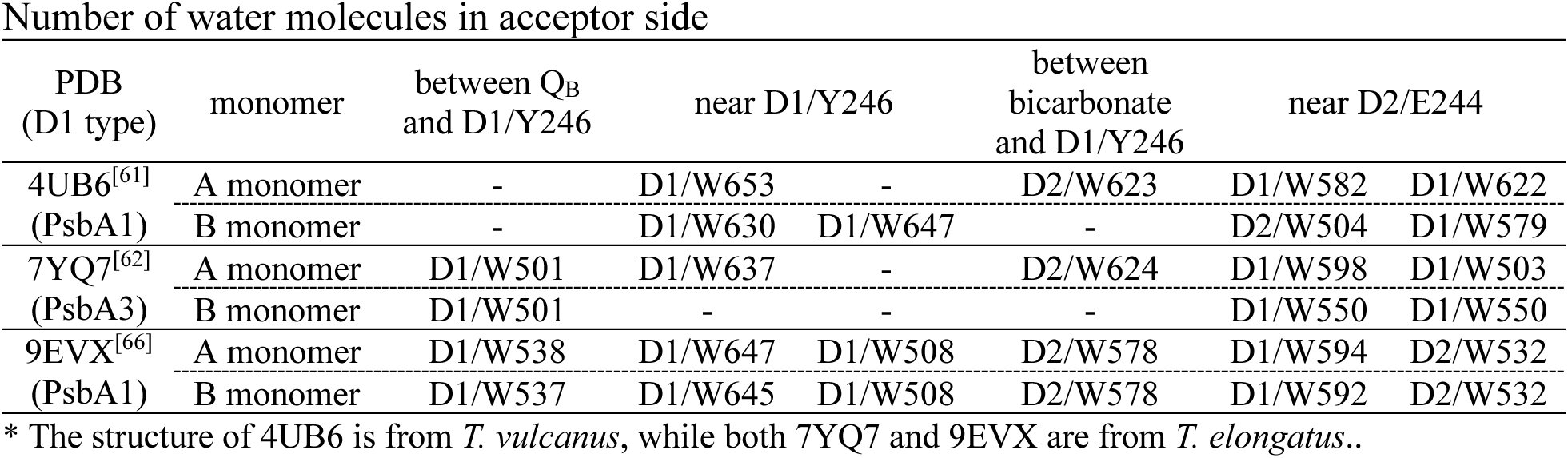
Number of water molecules in acceptor side

| PDB<br>(D1 type) | monomer | between Q<sub>B</sub><br>and D1/Y246 | near D1/Y246 |  | between<br>bicarbonate<br>and D1/Y246 | near D2/E244 |  |
| --- | --- | --- | --- | --- | --- | --- | --- |
| 4UB6<sup>[61]</sup> | A monomer | - | D1/W653 | - | D2/W623 | D1/W582 | D1/W622 |
| (PsbA1) | B monomer | - | D1/W630 | D1/W647 | - | D2/W504 | D1/W579 |
| 7YQ7<sup>[62]</sup> | A monomer | D1/W501 | D1/W637 | - | D2/W624 | D1/W598 | D1/W503 |
| (PsbA3) | B monomer | D1/W501 | - | - | - | D1/W550 | D1/W550 |
| 9EVX<sup>[66]</sup> | A monomer | D1/W538 | D1/W647 | D1/W508 | D2/W578 | D1/W594 | D2/W532 |
| (PsbA1) | B monomer | D1/W537 | D1/W645 | D1/W508 | D2/W578 | D1/W592 | D2/W532 |

\* The structure of 4UB6 is from *T. vulcanus*, while both 7YQ7 and 9EVX are from *T. elongatus*..

